# TREK1 upregulation is an endogenous mechanism delaying cognitive decline in Alzheimer’s Disease

**DOI:** 10.1101/2025.10.16.682816

**Authors:** Tuhina Mitra, Ranjit Bhoi, Triparna Chakraborty, Asmit Moharana, Vaishnav Manoj, Hiral Rawal, Swagata Ghatak

**Author notes:** Corresponding author: Dr. Swagata Ghatak. These authors have contributed equally (Tuhina Mitra, Ranjit Bhoi).

## Abstract

Alzheimer’s Disease (AD) is marked by early hippocampal and neocortical accumulation of amyloid-beta 42 oligomers (Aβ42o), driving neuronal hyperactivity and synaptic dysfunction years before symptom onset. While two-pore domain leak potassium channels like TREK1 are crucial for maintaining the resting membrane potential of neurons and shaping their excitability profile, their role in major neurodegenerative disorders like AD remains unknown. Here, we discover an activity-dependent upregulation of TREK1 in AD transgenic mice (3xTg and APP/PS1) and cultured hippocampal/cortical neurons, triggered by Aβ42o -induced hyperactivity. Mechanistically, we show that increased intracellular calcium activates adenylate cyclase 1/8 (AC1/8), initiating a cAMP-PKA signaling cascade that enhances the expression of chromatin regulator CTCF. This increased CTCF in turn enhances the expression of TREK1 both *in vitro* and in AD transgenic mice. Through a combination of calcium imaging, patch-clamp electrophysiology, immunostaining, and cognitive assays in a hippocampal TREK1 knockdown AD model, we establish that the pathology-associated upregulation of TREK1 constitutes a critical homeostatic brake on neuronal hyperexcitability. This neuronal response is essential for limiting Aβ42o-mediated synaptic pathology and delaying cognitive decline. Our study identifies a multi-step signaling cascade triggered by Aβ42o leading to upregulation of TREK1 that functions as an essential compensatory mechanism for neuronal survival in early AD. This study deciphers a cellular mechanism responsible for the preclinical phase of AD characterized by silent buildup of pathology.

## Introduction

AD is a leading cause of dementia, characterized by progressive cognitive decline. Emerging evidence indicates that AD patients exhibit heightened neuronal network hyperactivity, which contributes to epileptiform spikes^1,2^. Both familial (FAD) and sporadic (SAD) forms of AD display neuronal hyperexcitability with onset in early stages of the disease^3,4^. Functional MRI studies reveal hyperactivation in the right anterior hippocampus of early-stage FAD patients^5^. Additionally, AD models—including transgenic mice, human iPSC-derived neurons, and cerebral organoids—recapitulate this network hyperactivity ^2, 6–8^.

One of the reasons for this hyperactivity is the pathological accumulation and aggregation of amyloid β ^9,10^ (Aβ). The disease process begins with the buildup of soluble amyloid-beta oligomers (Aβo) in the neocortex and hippocampus, occurring at least a decade before clinical symptoms emerge and preceding the formation of insoluble Aβ plaques^11^. These Aβo disrupt neuronal function by inducing hyperactivity, elevating intracellular calcium levels, and ultimately driving synaptic and neuronal degeneration^9, 12, 13^. Among the Aβ peptides, Aβ42 and Aβ40 are the primary isoforms that oligomerize and accumulate in AD^8, 14^. Despite being only two residues longer than Aβ40, Aβ42 exhibits significantly higher toxicity due to its enhanced aggregation propensity^15, 16^. Notably, Aβ42o have been shown to trigger pathological calcium influx and upregulate the sodium channel Nav1.6, exacerbating neuronal hyperexcitability^17^. Aβ42o-mediated suppression of Kv1.1 activity has been linked to neuronal hyperactivity in AD^18,19^. Further, Aβ42 decreased synaptic inhibition via internalization of GABAA receptors contributing to neuronal network hyperexcitability^20^. Aβ42o increased the NR2B-containing NMDAR density in neuronal dendrites and increased intracellular Ca^2+^ levels^21^.

Neuronal activity is tightly regulated by a diverse array of ion channels, among which two-pore domain leak potassium channels (K2P) play a critical role in maintaining resting membrane potential and modulating excitability^22–24^. Beyond neurons, astrocytic K2P contributes to glutamate clearance and ionic homeostasis, particularly under ischemic stress^25–27^. Among the various K2P channels, TREK1 subfamily of K2P channels has emerged as a key neuroprotective player in stroke and epilepsy where neuronal hyperactivity is a key feature^28^. TREK1 is one of most studied K2P and is highly expressed in several regions of the brain known to be affected by AD like the hippocampus, and the cortex^29^. Intriguingly, recent work in the SAMP8 mouse model (a genetically accelerated aging model) revealed a significant upregulation of TREK1 expression in young animals compared to controls^30^. This raised two key unresolved questions in the context of AD-1) if and how TREK1 expression is altered in AD? 2) how TREK1 modulation, in turn, influences neuronal viability and network hyperexcitability, a hallmark of early AD? 3) whether TREK1 modulation affects cognitive function which is known to deteriorate as AD progresses?

Using immunostaining, calcium imaging, patch clamp electrophysiology and cognitive assays we discover a novel multistep signalling cascade triggered by Aβ-related hyperactivity that upregulates TREK1 expression. Surprisingly, we found neuronal hyperexcitability itself increases TREK1 expression which acts like negative feedback to decrease excitotoxicity. This mitigates AD pathology and improves cognitive function.

## Results

### Increased TREK1 expression in AD transgenic mice

We observed a significant increase in TREK1 expression in hippocampal and cortical regions of 3-month-old APP/PS1 (Fig. 1a-e and k) and 3xTg mice (Fig. 1f-k and Supplementary Fig. 1a). The increase in TREK1 expression was ∼1.5-2 fold by immunohistochemistry and ∼3-4 fold by immunoblotting in APP/PS1 mice (Fig. 1b,c and e) whereas in 3xTg the increase was ∼1.5 fold by both the immunostaining techniques (Fig. 1g, h and j). These models were selected for their distinct AD-relevant pathological features: APP/PS1 mice^31^ (expressing Swedish APP + PSEN1dE9) show age-dependent Aβ42o accumulation, while 3xTg-AD mice (Swedish APP + P301L tau + M146V PSEN1) develop both Aβ plaques and neurofibrillary tangles ^32^. Both models exhibited neuronal hyperactivity manifested as increased calcium transients in acute hippocampal slices (Supplementary Fig. 1b-e and Supplementary Video 9 and 10) and early AD pathology, including elevated Aβ levels (Supplementary Fig. 1f-i), consistent with established phenotypes. We studied pathophysiology in the hippocampus and cortex as they are most prone to AD^33, 34^.

**Fig. 1.**
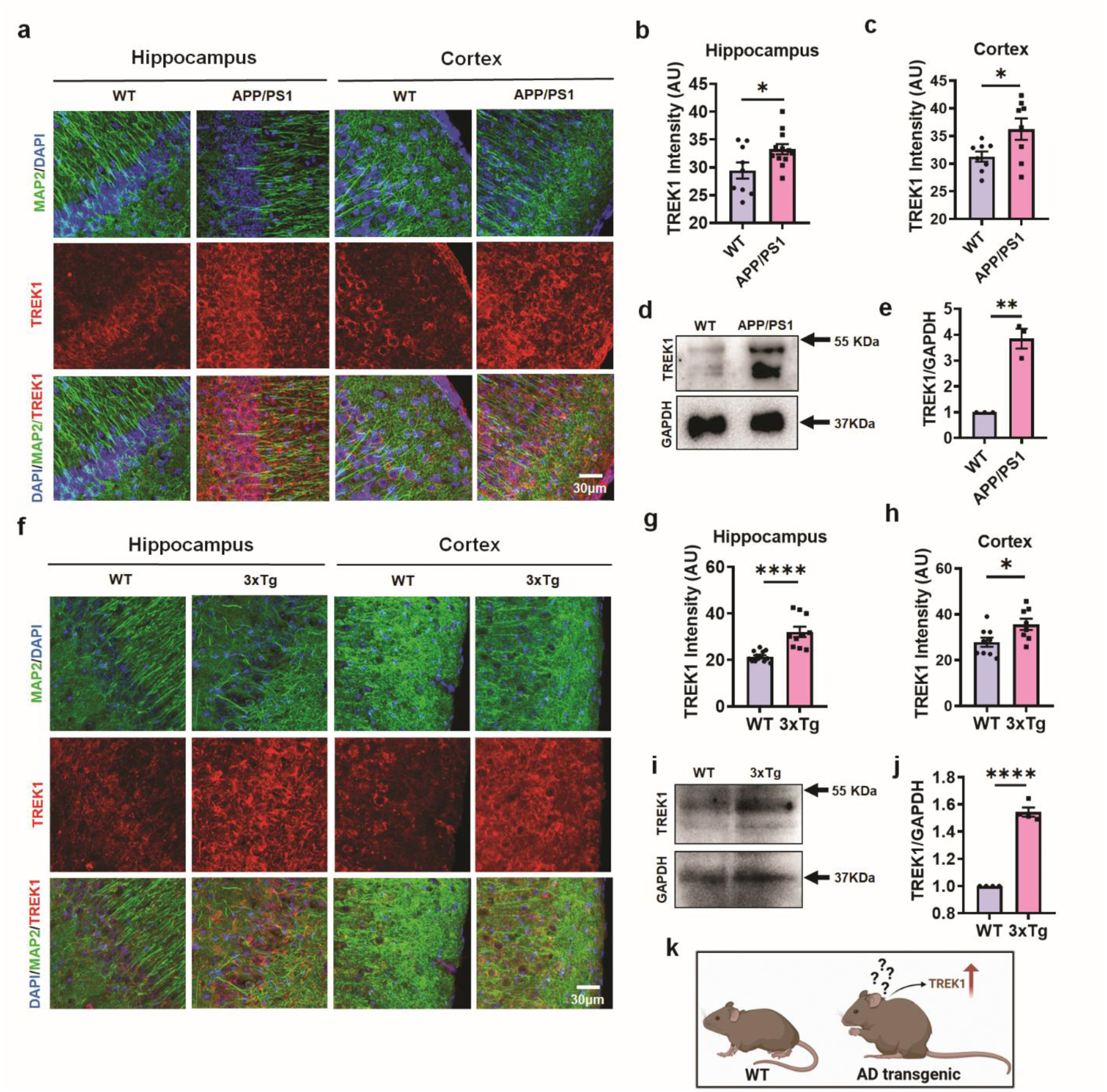
Upregulation of TREK1 in transgenic mouse models of Alzheimer’s disease. a, Representative immunofluorescence images showing elevated TREK1 expression in the hippocampus and cortex of APP/PS1 mice compared to age-matched wild-type controls. b, Quantification of TREK1 fluorescence intensity in the APP/PS1 mice hippocampus (n=9-12 sections; *P < 0.05; two-tailed unpaired t-test). c, Quantification of TREK1 fluorescence intensity in the APP/PS1 mice cortex (n=8 sections; *P < 0.05; two-tailed unpaired t-test). d, Western blot of brain cortex lysates showing increased TREK1 protein abundance in APP/PS1 mice. e, Densitometric analysis and quantification of TREK1 protein levels normalized to GAPDH (n=3; **P < 0.01; two-tailed unpaired t-test). f, Representative immunofluorescence images showing increased TREK1 expression in the hippocampus and cortex of 3xTg mice compared to wild-type controls. g, Quantification of TREK1 fluorescence intensity in the 3xTg mice hippocampus (n=10-14 sections; ****P < 0.0001; two-tailed unpaired t-test). h, Quantification of TREK1 fluorescence intensity in the 3xTg mice cortex (n=8-9 sections; *P < 0.05; two-tailed unpaired t-test). i, Western blot analysis shows increased TREK1 protein abundance in 3xTg mouse brain homogenates. j, Densitometric analysis and quantification of TREK1 protein levels normalized to GAPDH in 3xTg mice (n=3; ****P < 0.0001; two-tailed unpaired t-test). k, Schematic representation showing increased TREK1 expression in AD transgenic mice. Data are presented as mean ± SEM. 3–4 mice per group were used.

Notably, in vitro treatment with 5 µM Aβ42o increased TREK1 expression significantly in primary hippocampal and cortical neurons after 24h by immunocytochemistry (∼1.25 fold) and immunoblotting (∼2 fold; Fig. 2a-d and Supplementary Fig. 2a, b). Interestingly, equimolar Aβ42 monomers and Aβ40 oligomers could not increase TREK1 levels significantly as observed by immunocytochemistry (Fig. 2a, b). This effect was replicated in vivo through intrahippocampal Aβ42o injection in WT mice (C57BL/6), with immunostaining analysis of hippocampal sections showing significant TREK1 upregulation (Fig. 2e-f). However, when Aβ40 oligomers were injected into the brain of WT mice, there was no increase in TREK1 expression (Supplementary Fig. 2c, d). The results suggested that Aβ42o accumulation in early AD led to the upregulation of TREK1 in mice brains.

**Fig. 2.**
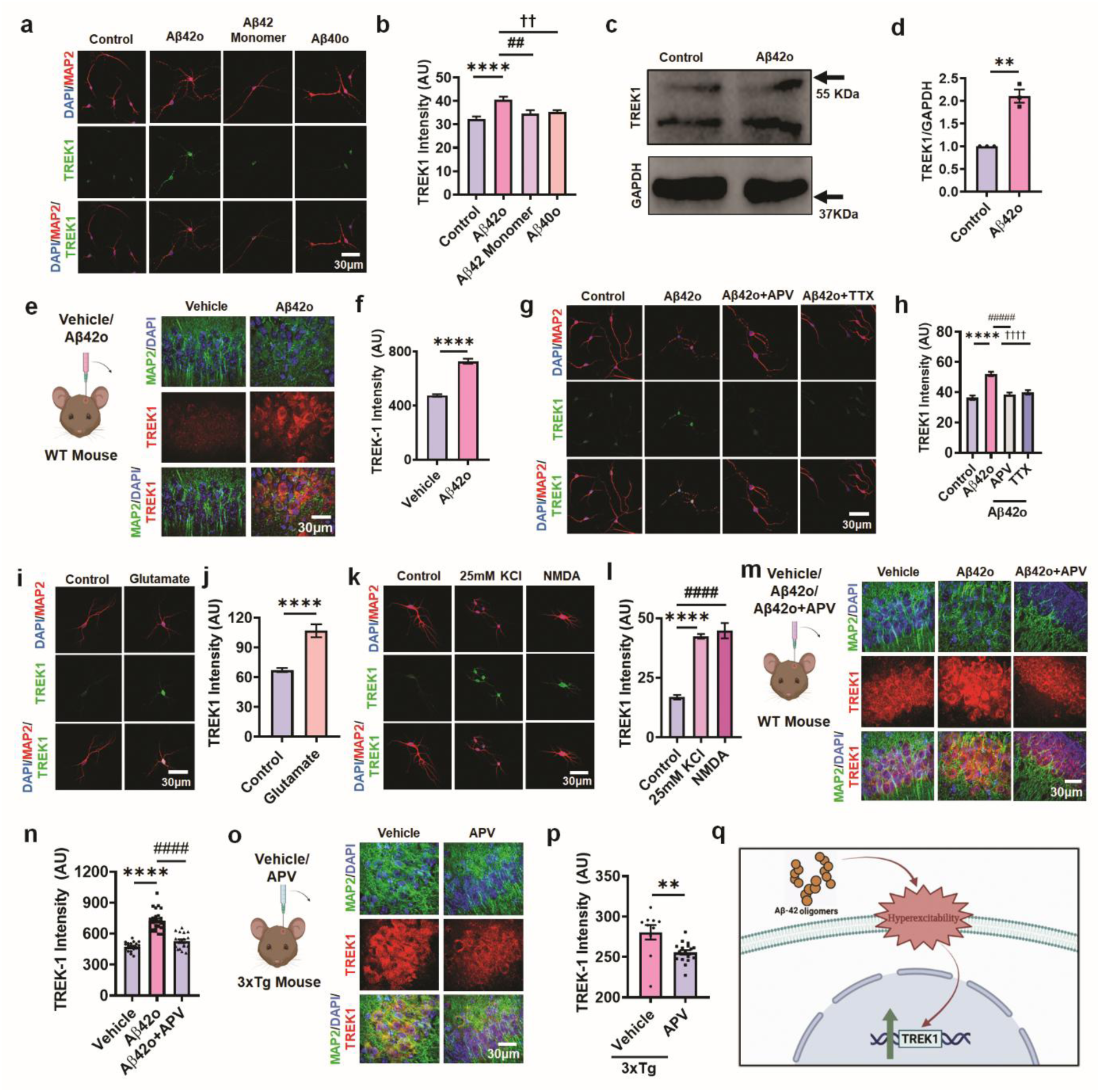
Aβ42o related neuronal hyperactivity induces TREK1 upregulation in primary neurons and the mouse brain. a, Representative immunofluorescence images showing increased TREK1 expression in primary neurons treated with Aβ42o compared with Aβ42 monomers, Aβ42o, and control. b Quantification of TREK1 fluorescence intensity upon treatment with Aβ42o, Aβ42 monomer, and Aβ40o compared with control (n=46-76 cells; ****p < 0.0001, ##p < 0.001, ††p < 0.001, one-way ANOVA followed by Šidák’s multiple comparisons test). c, Western blot analysis of primary neuron–astrocyte co-cultures confirming increased TREK1 levels 42o treated neurons compared with controls. d, Densitometric analysis and quantification of TREK1 protein levels normalized to GAPDH in Aβ42o treated co-cultures (n=3; **p < 0.01, unpaired t-test). e, Hippocampal sections from C57BL/6 wild-type mice 24 hours after intrahippocampal injection of Aβ42o showing increased TREK1 expression compared with vehicle-injected controls. f, Quantification of TREK1 expression from (e) (n=21-27 sections; ****p < 0.0001, unpaired t-test). g, Representative images showing that co-treatment of Aβ42o with APV or TTX reduces Aβ42o-induced TREK1 upregulation in primary cultures. h, Quantification of TREK1 fluorescence intensity in APV/TTX treated primary cultures (n=88-154 cells; ****p < 0.0001, ####p < 0.0001, ††††p < 0.0001, one-way ANOVA with Šidák’s test). i, Exposure to glutamate mimics Aβ42o-induced TREK1 upregulation. j, Quantification of TREK1 fluorescence intensity following glutamate treatment (n=36-39 cells; **** p < 0.0001, unpaired t-test). k, Treatment with NMDA or KCl to induce neuronal hyperexcitability increases TREK1 levels. l, Quantification of TREK1 fluorescence intensity following KCl and NMDA treatment (n=25-79 cells; ****p < 0.0001, ####p < 0.0001, one-way ANOVA with Šidák’s test). m, Representative immunofluorescence images of brain sections from Aβ42o injected mice showing elevated TREK1 expression in the hippocampus, which is attenuated by co-injection with APV. n, Quantification of hippocampal TREK1 intensity after injecting Aβ42o and/or APV (n=18-27 sections; ****p < 0.0001, one-way ANOVA followed by Šidák’s multiple comparisons test). o, Representative images of hippocampal sections from 3xTg mice treated with APV or vehicle showing decreased TREK1 expression following NMDA receptor blockade. p, Quantification of TREK1 intensity following APV treatment in 3xTg mice (n=10-18 sections; **p < 0.01, unpaired t-test). q, Schematic representation illustrating that Aβ42o induced neuronal hyperexcitability drives increased TREK1 expression. Data are presented as mean ± SEM. 3-5 independent cultures or animals per group were used.

### Aβ42o-induced neuronal hyperexcitability drives TREK1 upregulation

In line with previous literature, our calcium imaging studies demonstrate that Aβ42o (5 µM) cause neuronal hyperactivity indicated by an increase in neuronal calcium transient frequency in primary neuron astrocyte co-cultures^35, 36^ (Supplementary Fig. 2e, f and Supplementary Video 1, 2). This led to a subsequent increase in TREK1 expression through an activity-dependent mechanism (Fig. 2g-q). The Aβ42o-mediated hyperexcitability is decreased with 50 µM APV (2-Aminophosphonovaleric acid, NMDA receptor antagonist) or 1 µM TTX (Tetrodotoxin, voltage-gated sodium channel blocker) (Supplementary Fig. 2e, f). Thereafter, we investigated whether suppressing neuronal hyperactivity could restore normal TREK1 expression. We observed that treating neuron astrocyte co-cultures with 50 µM APV or 1 µM TTX indeed decreased TREK1 expression similar to that of control levels as quantified by immunostaining (Fig. 2g, h). We further confirmed the activity-dependence of TREK1 expression through three independent neuronal hyperexcitability paradigms in primary neuron astrocyte co-cultures: treatment with 100 µM glutamate, 25 mM KCl, or 100 µM NMDA. Interestingly all the experimental approaches used to induce hyperactivity replicated the increase in TREK1 expression (Fig. 2i-l and Supplementary Fig. 2g). Importantly, in vivo validation showed that intrahippocampal co-injection of Aβ42o with APV in WT mice prevented the TREK1 upregulation observed with Aβ42o injection alone, as demonstrated by immunohistochemistry of hippocampal sections (Fig. 2m, n). Additionally, we injected APV in the hippocampus of 3xTg mice and found a decrease in TREK1 levels suggesting that neuronal hyperactivity was able to induce TREK1 expression (Fig. 2o, p). These complementary in vitro and in vivo findings demonstrate that Aβ42o-induced upregulation of TREK1 is driven by neuronal hyperexcitability, revealing a novel activity-dependent mechanism regulating TREK1 expression.

### Calcium-dependent activation of adenylyl cyclase 1/8 upregulates TREK1 expression

To elucidate the signaling pathway underlying activity-dependent TREK1 expression, we investigated the role of intracellular calcium. Since neuronal hyperactivity elevates intracellular calcium^37^, we first assessed the effect of calcium chelation on TREK1 expression. Treatment with BAPTA-AM, a calcium chelator, reduced neuronal calcium transients even in the presence of Aβ42o in primary neurons in calcium imaging studies (Supplementary Fig. 3a, b). BAPTA-AM significantly decreased TREK1 expression in primary neurons exposed to Aβ42o by immunocytochemistry confirming that increased intracellular calcium was driving the expression of TREK1 (Fig. 3a, b and Supplementary Fig. 3c).

**Fig. 3.**
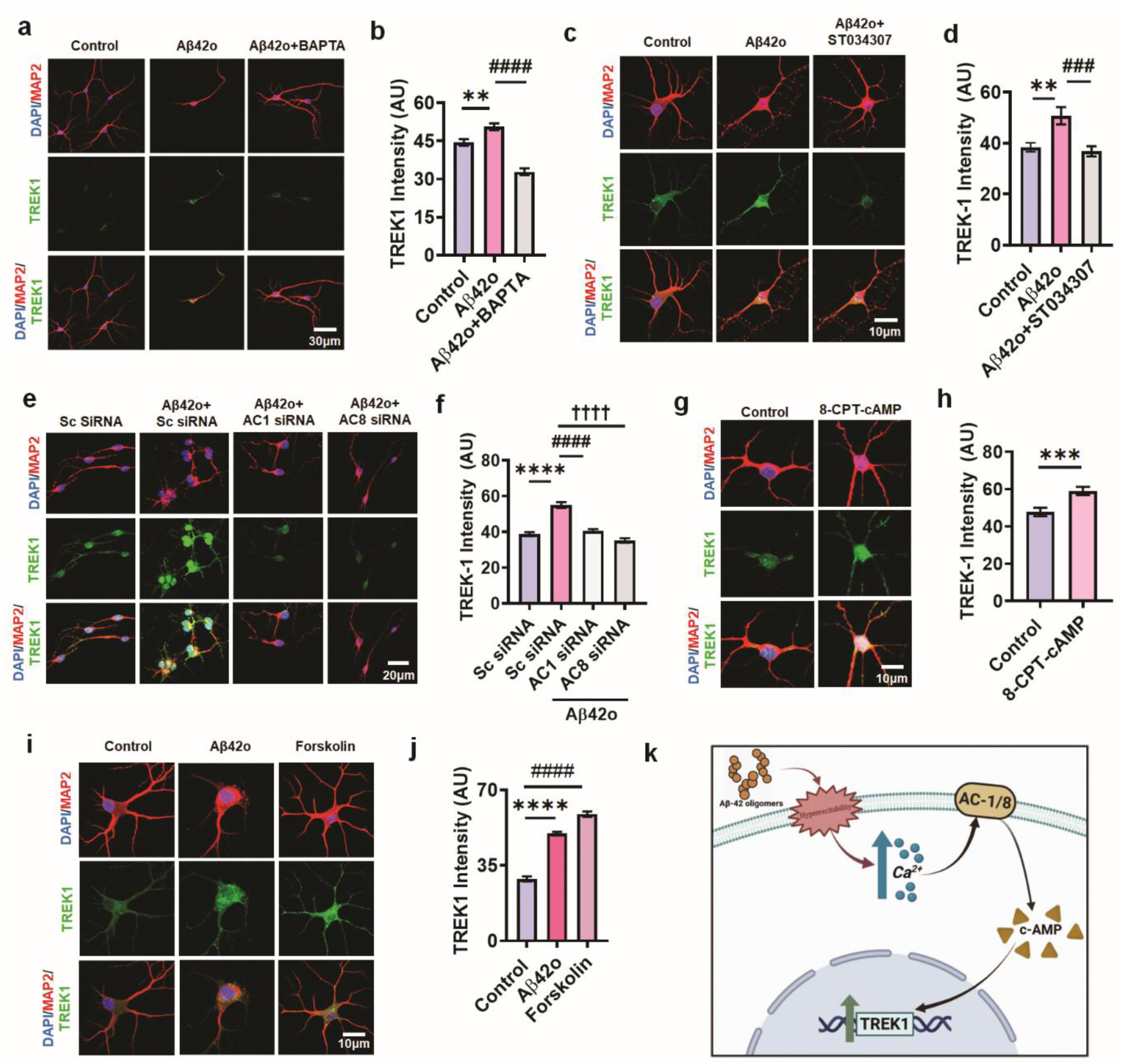
Aβ42o induce TREK1 upregulation in primary neurons via AC1/AC8–cAMP signaling pathway triggered by increased intracellular calcium. a, Representative immunofluorescence images showing increased TREK1 expression upon Aβo treatment, which is markedly reduced by co-treatment with the calcium chelator BAPTA-AM. b, Quantification of TREK1 fluorescence intensity following Aβ42o and/or BAPTA-AM treatment (n=61-90 Cells; **p < 0.01, ####p < 0.0001; one-way ANOVA with Šidák’s test). c, Representative images showing that TREK1 increase induced by Aβ42o is attenuated by the AC1 inhibitor ST034307. d, Quantification of TREK1 fluorescence intensity following Aβ42o and/or ST034307 treatment (n=23-31 cells; **p < 0.01, ###p < 0.001; one-way ANOVA with Šidák’s test). e, Representative images showing TREK1 expression is decreased in Aβ42o-treated neurons transfected with AC1-specific or AC8-specific siRNA, but not with scrambled (Sc) siRNA. f, Quantification of TREK1 fluorescence intensity following Aβ42o treatment with AC1 or AC8 knockdown (n=52-93 cells; ****p < 0.0001, ####p < 0.0001, ††††p < 0.0001; one-way ANOVA with Šidák’s test). g, Treatment with the cAMP analog 8-CPT-cAMP increases TREK1 expression in primary neurons. h, Quantification of TREK1 fluorescence intensity following 8-CPT-cAMP treatment (n=57-62 cells; ***p < 0.001, unpaired t-test). i, Treatment with forskolin, a cAMP activator, mimics Aβ42o by increasing TREK1 expression in primary neurons. j, Quantification of TREK1 fluorescence intensity following forskolin treatment (n=146-242 cells; ****p < 0.0001, ####p < 0.0001; one-way ANOVA with Šidák’s test). k, Schematic representation illustrating that Aβ42o-induced TREK1 upregulation is mediated by calcium influx via the AC1/AC8–cAMP signaling pathway. Data are presented as mean ± SEM. 3-5 independent cultures per group were used.

Previous studies indicate that elevated calcium enhances the activity of specific adenylyl cyclase (AC) isoforms. Among the known AC isoforms, AC1 and AC8 are uniquely regulated by calcium and calmodulin, enabling them to directly convert activity-dependent calcium rise into cAMP production^38^. This coupling can further amplify signaling through downstream effectors such as protein kinase A (PKA)^39^. Biochemical and genetic studies have established that AC1 and AC8 are the predominant calcium-sensitive ACs in the central nervous system^40^, positioning them as critical mediators of activity-regulated cAMP signaling in neurons. To determine their involvement, we pharmacologically inhibited AC1 using ST034307 (ST) which abolished Aβ42o-induced TREK1 upregulation in primary neurons, as demonstrated by immunocytochemistry (Fig. 3c, d). Further, we found a similar decrease in TREK1 expression upon treatment of primary neuron astrocyte co-cultures with AC1 siRNA and AC8 siRNA in addition to Aβ42o suggesting that both AC1 and AC8 play an important role in Aβ42o-induced TREK1 upregulation (Fig. 3e, f).

Since AC1/AC8 activation elevates intracellular cAMP, we investigated whether cAMP directly modulates TREK1 levels in neurons. We applied the membrane-permeable cAMP analog 8-CPT-cAMP on primary neuron astrocyte co-cultures and observed a similar enhancement of TREK1 levels demonstrating that cAMP elevation alone is sufficient to upregulate TREK1 (Fig. 3g, h). To confirm this, we treated the primary neurons with forskolin, an AC activator and found significant increase in TREK1 expression, further validating cAMP as the key downstream effector of AC1/AC8 signaling (Fig. 3i, j and Supplementary Fig. 3c). This is in line with previous studies which have shown that both 8CPT-cAMP and forskolin increased the expression of TREK1 mRNA in bovine adrenal zona fasciculata and rat astrocytes respectively^26, 39^. Together, these findings establish that Aβ42o related hyperactivity trigger a calcium-AC1/AC8-cAMP cascade that drives TREK1 overexpression in neurons (Fig. 3k).

### cAMP drives TREK1 upregulation via the PKA-CTCF signaling

Since PKA is one of the major downstream effectors of cAMP and cAMP has been shown to directly activate PKA^41^, we hypothesized that PKA could be causing the cAMP-mediated increase in TREK1 expression. Therefore, we inhibited PKA using pharmacological blockers like KT5720 and H89 which abolished the increase in TREK1 levels in primary neurons detected by immunocytochemistry (Fig. 4 a-d and Supplementary Fig. 4a).

**Fig. 4.**
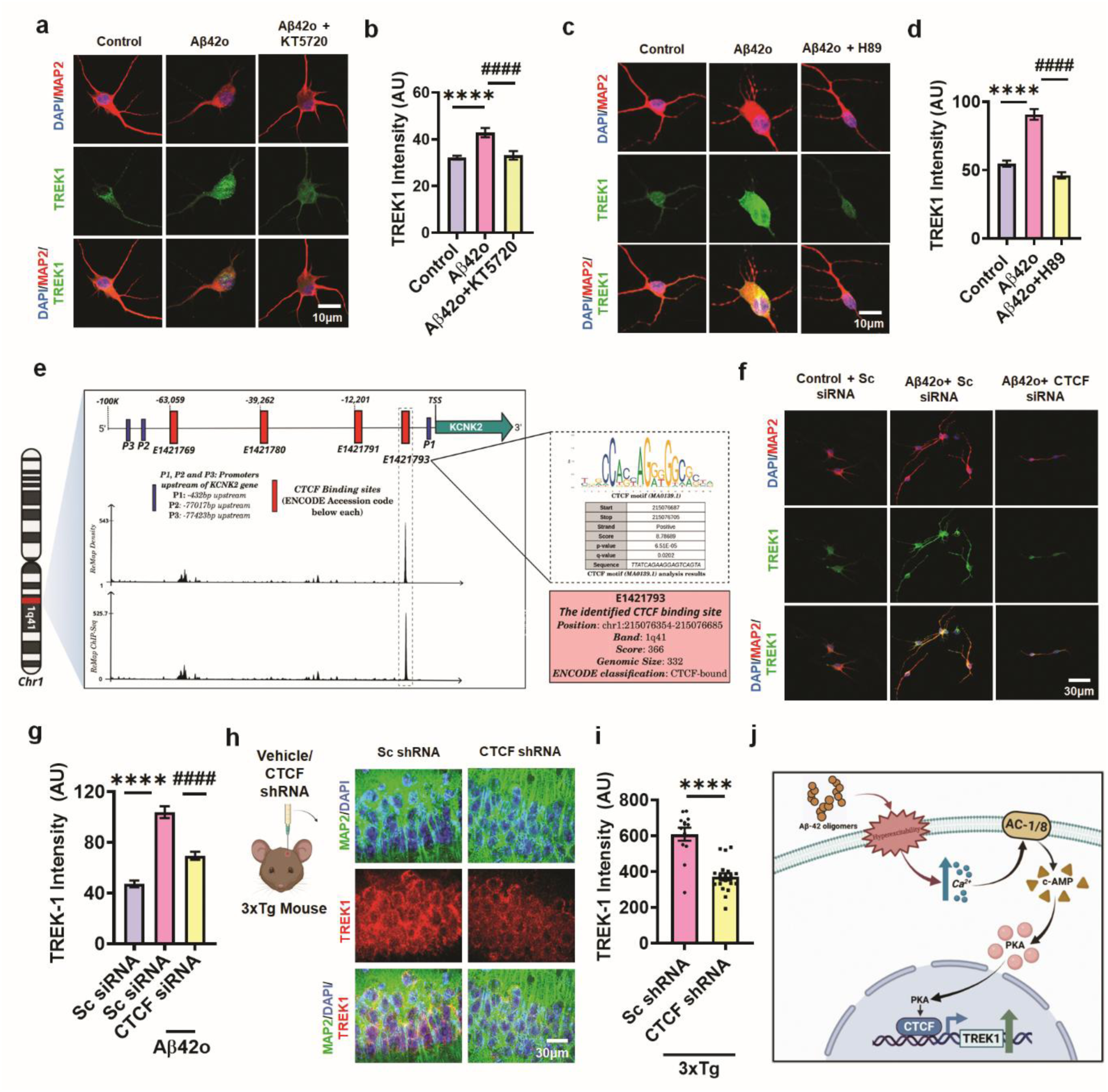
The PKA–CTCF signaling axis regulates Aβ42o-induced TREK1 expression. a, Representative immunofluorescence images showing increased TREK1 expression in Aβ42o-treated neurons, which is decreased upon co-treatment with the PKA inhibitor KT5720. b, Quantification of TREK1 fluorescence intensity following Aβ42o and/or KT5720 treatment (n = 52-70 cells; ****p < 0.0001, ####p < 0.0001; one-way ANOVA with Šidák’s test). c, Representative images showing that TREK1 increase induced by Aβ42o is attenuated by another PKA inhibitor H89. d, Quantification of TREK1 fluorescence intensity following Aβ42o and/or H89 treatment (n = 43–49 cells; **** p < 0.0001, ####p < 0.0001; one-way ANOVA with Šidák’s test). e, Schematic representation of the TREK1 locus on chromosome 1q41 showing predicted promoters (blue boxes; P1–P3) and ENCODE-annotated CTCF binding sites (red boxes; accession IDs indicated). Genomic positions are indicated relative to the transcription start site (TSS). ReMap ChIP-seq and density tracks demonstrate experimental support for the predicted binding sites, with peaks at site E1421793 located proximal to the promoter (P1). Motif analysis using JASPAR confirmed the presence of a consensus CTCF motif within this region (right panel), with associated FIMO statistics (score, p-value, and q-value). The identified site (highlighted in red) spans chr1:215076354–215076685 (band 1q41), has a genomic size of 332 bp, and is classified as “CTCF-bound” in ENCODE. f, Representative images showing decreased TREK1 expression in Aβ42o treated neurons transfected with CTCF-specific siRNA compared to scrambled (Sc) siRNA. g, Quantification of TREK1 fluorescence intensity following Aβ42o and/or CTCF knockdown (n = 37–76 cells; **** p < 0.0001, #### p < 0.0001; one-way ANOVA with Šidák’s test). h, Representative images showing decreased TREK1 expression in hippocampal neurons of 3xTg mice following intrahippocampal injection of CTCF shRNA lentivirus compared to control. i, Quantification of TREK1 fluorescence intensity in 3xTg mice after CTCF knockdown (n = 13–19 sections; ****p < 0.0001; unpaired t-test). j, Schematic representation of the PKA–CTCF signaling axis in regulating Aβ42-induced TREK1 expression. Data are presented as mean ± SEM. 3-5 independent cultures or animals per group were used.

However, it was not clear how PKA was causing this increase in expression of TREK1. We went on to check enhancer and promoter regions present in TREK1 cistron. We found from a cistronic analysis done on signalingpathways.org that CTCF binding site was commonly found associated with the TREK1 gene. We computationally identified potential CTCF binding sites in the TREK1 (KCNK2) regulatory region by analyzing its genomic sequence (gene body + 100 kb upstream of transcription start site) from the UCSC Genome Browser (hg38/GENCODE v48; Fig. 4e). Using MEME Suite’s FIMO tool, we detected high-confidence CTCF motifs (JASPAR: MA0139.1; p < 0.0001) and mapped these sites relative to known promoters (validated via Eukaryotic Promoter Database). To ensure reliable identification of functional CTCF regulatory elements in the TREK1 locus we used a workflow integrating ENCODE cCREs, JASPAR predictions, and ReMap ChIP-seq data (Fig. 4e and Supplementary Fig. 5a). Additionally, a study enlisting genes that were down-regulated in the CTCF-cKO mice reported KCNK2 to be downregulated in these mice^42^. Interestingly, PKA has been shown to upregulate CTCF transcript levels^43^ and on application of PKA blocker H89 in primary neuron astrocyte co-cultures, we found a decrease in CTCF levels (Supplementary Fig. 4b, c). Based on these findings we went on to knockdown CTCF with siRNA in primary neurons (Supplementary Fig. 4d, e) and found that Aβ42o were unable to increase TREK1 expression (Fig. 4f, g and Supplementary Fig. 4f). These results indicated that CTCF is involved in the Aβ42o-mediated change in expression of TREK1. Further, to establish the *in vivo* relevance of our findings, we performed targeted CTCF knockdown in 3xTg-AD mice through stereotaxic delivery of lentiviral construct bearing CTCF shRNA to the hippocampus, confirmed by immunohistochemistry (Supplementary Fig. 4g, h). CTCF knockdown in 3xTg mice hippocampus led to a significant reduction in TREK1 expression as determined by immunohistochemistry (Fig. 4h, i). These experiments led us to conclude that cAMP was acting via PKA-CTCF pathway to increase TREK1 expression (Fig. 4j)

### Increased TREK1 expression decreases pathological neuronal activity, limits excitatory/inhibitory (E/I) imbalance

Since Aβ42o-related neuronal hyperactivity was able to increase TREK1 expression via a Ca^2+^-AC1/8-cAMP-PKA-CTCF pathway, we asked what could be the reason for such upregulation of TREK1 expression in neurons? To assess TREK1’s role, we pharmacologically inhibited TREK1 (using specific inhibitors, spadin) or knocked it down via siRNA in neurons exposed to Aβ42o (Supplementary fig. 6a). Both approaches markedly increased neuronal hyperexcitability, as evidenced by elevated calcium event frequencies measured using Fluo4 AM calcium imaging dye (Fig. 5a, b, e and f and Supplementary Video1, 2, 3, 5, 6 and 7) and membrane potential spikes measured using FluoVolt compared to Aβ42o treatment alone (Fig. 5c, d). Conversely, TREK1 activation pharmacologically (using BL1249) or overexpression (Supplementary fig. 6b) significantly reduced calcium event frequencies, counteracting Aβ42o-induced hyperexcitability (Fig. 5a, b, e and g and Supplementary Video1, 2, 4, 5, 6 and 8). Further by patch clamp electrophysiology we observed that the spadin-treated neurons showed increased spontaneous action potential (AP) frequency (Fig. 5h, i) and depolarized resting membrane potential (RMP) compared to Aβ42o-treated neurons (Fig. 5j) indicating that TREK1 activity was important for limiting Aβ-induced hyperactivity and membrane depolarization in neurons. Similarly, excitatory postsynaptic current (EPSC) frequency was significantly increased upon inhibition of TREK1 by spadin in Aβ42o-treated neurons compared to Aβ42o treatment alone (Fig. 5k, l). The EPSC amplitude remained unchanged (Fig. 5m). Inhibitory postsynaptic current (IPSC) frequency was significantly decreased upon inhibition of TREK1 by spadin in Aβ42o treated neurons compared to Aβ42o treatment alone (Fig. 5n, o). The IPSC amplitude remained unchanged (Fig. 5p). This increase in EPSC frequency and decrease in IPSC frequency would lead to an increase in the E/I ratio upon TREK1 inhibition compared to Aβ42o treatment alone disrupting the E/I balance further. These findings suggest that TREK1 upregulation serves as a compensatory neuroprotective mechanism, dampening aberrant neuronal activity triggered by Aβ42o. To further confirm the effect of TREK1 on neuronal hyperactivity in AD, we injected 3xTg mice with lentiviral TREK1 shRNA (Supplementary Fig. 6c). Upon silencing TREK1, 3xTg mice showed increased calcium events in ex-vivo hippocampal acute slices compared to vehicle injected 3xTg mice (Fig. 5q-s and Supplementary Video 11 and 12). Therefore, we concluded that TREK1 is important in limiting the existing pathological neuronal excitability in 3xTg mice.

**Fig. 5.**
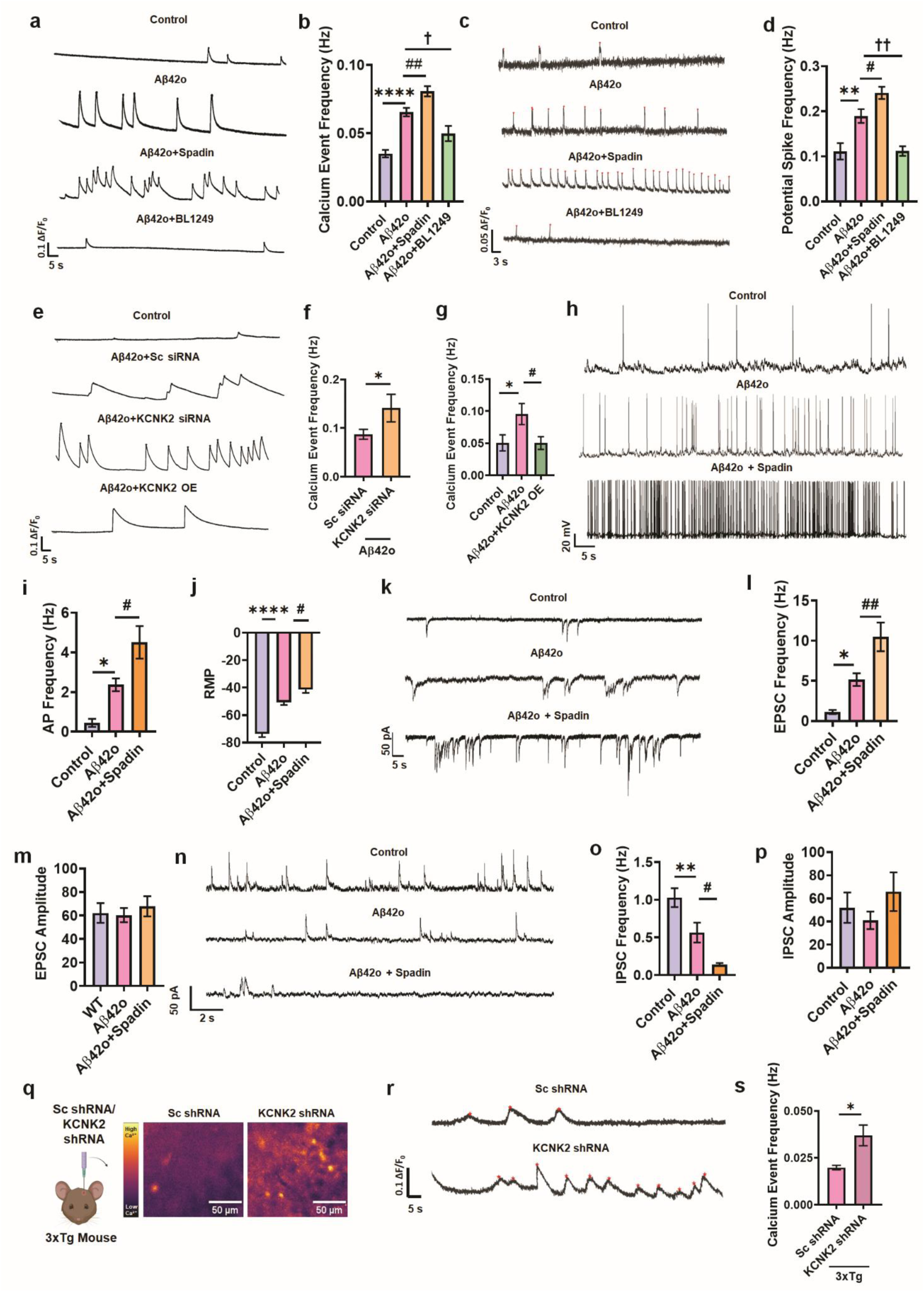
TREK1 controls Aβ42o-induced aberrant neuronal activity and E/I ratio. a, Representative calcium imaging traces from control and Aβ42o treated neurons showing that Aβ42o increases spontaneous calcium transient frequency. This hyperexcitability is further enhanced by the TREK1 inhibitor spadin and suppressed by the TREK1 activator BL-1249. b, Quantification of calcium event frequency upon treatment with Aβ42o/spadin/BL-1249 (n = 512–1002 cells; ****p < 0.0001, ##p < 0.01, †p < 0.05; one-way ANOVA with Šidák’s test). c, Representative FluoVolt traces measuring membrane potential fluctuations manifest enhanced neuronal activity in the presence of TREK1 inhibitor spadin and suppressed neuronal activity by the TREK1 activator BL-1249 compared to Aβ42o treatment alone. d, Quantification of potential spike frequency upon treatment with Aβ42o/spadin/BL-1249 (n = 31–105 cells; **p < 0.01, #p < 0.05, ††p < 0.01; one-way ANOVA with Šidák’s test). e, Representative calcium traces from neurons treated with Aβ42o along with scrambled (Sc) siRNA, KCNK2 siRNA, or a KCNK2 overexpression (OE) construct. f, Quantification of calcium event frequency upon knocking down KCNK2 in the presence of Aβ42o treatment (n = 49–67 cells; *p < 0.05; unpaired t-test). g, Quantification of calcium event frequency upon overexpressing KCNK2 in the presence of Aβ42o treatment (n = 37–48 cells; *p < 0.05, #p < 0.05; one-way ANOVA with Šidák’s test). h, Representative patch-clamp recordings of action potentials in control, Aβ42o, and Aβ42o + spadin treated neurons showing exacerbated action potential firing with TREK1 blockade. i, Quantification of action potential frequency following Aβ42o and/or spadin treatment (n = 17 cells; *p < 0.05, #p < 0.05; one-way ANOVA with Šidák’s test). j, Resting membrane potential (RMP) is more depolarized in Aβ42o treated neurons compared to control, and further depolarizes in presence of spadin with Aβ42o (n = 19 cells; ****p < 0.0001, #p < 0.05; one-way ANOVA with Šidák’s test). k, Representative traces showing excitatory postsynaptic current (EPSC) frequency is increased in neurons treated with Aβ42o + spadin compared to Aβ42o alone. l, Quantification of EPSC frequency (n = 19 cells; *p < 0.05, ##p < 0.01; one-way ANOVA with Šidák’s test). m, Quantification of EPSC amplitude (n = 19 cells). n, Representative traces showing Inhibitory postsynaptic current (IPSC) frequency is decreased in neurons treated with Aβ42o+spadin compared to Aβ42o alone. o, Quantification of IPSC frequency (n = 10 cells; **p < 0.01, #p < 0.05; one-way ANOVA with Šidák’s test). p, Quantification of IPSC amplitude (n = 10 cells). q, Representative ex vivo calcium imaging heat map from hippocampal slices of 3xTg mice injected with TREK1 shRNA lentivirus showing elevated calcium activity compared to sc shRNA-injected mice. r, Representative calcium imaging traces demonstrating increased calcium transient frequency following TREK1 knockdown. s, Quantification of calcium event frequency in TREK1 knockdown mice compared to sc shRNA-injected mice (n = 13–27 cells; *p < 0.05; unpaired t-test). Data are presented as mean ± SEM from 3-5 independent cultures.

Since neuronal activity is directly linked to the number of synapses we quantified the total synapses by labelling synapsin-1 in hippocampal sections of 3xTg mice. We found decreased synapsin-1 expression in 3xTg mice compared with WT mice, reflecting a reduction in total synapses (Supplementary Fig. 7a, b). Further, to understand the effect of upregulated TREK1 on the E/I ratio of 3xTg mice hippocampus, we stained the neurons for VGLUT1 (which labels the excitatory presynapse) and VGAT (which labels the inhibitory presynapse). In line with previous literature, we found an increase in excitatory synapses and a decrease in inhibitory synapses in 3xTg mice, resulting in an increase in E/I ratio compared to WT^44^ (Supplementary Fig. 7c-g). To investigate the role of TREK1 in 3xTg Alzheimer’s disease (AD) mice, we injected TREK1 shRNA lentiviral particles into the hippocampus and compared the results to vehicle-injected controls. Silencing TREK1 led to a significant increase in excitatory synapses, as indicated by VGLUT1 staining, and a significant decrease in inhibitory synapses, as shown by VGAT staining (Fig. 6a–d). Consequently, the E/I ratio was elevated by approximately 1.5-fold in TREK1 knockdown mice compared to controls (Fig. 6e). These findings suggest that TREK1 silencing further disrupts the E/I balance in the hippocampus of AD mice, emphasizing the importance of TREK1 in maintaining normal neuronal activity and E/I ratio.

**Fig. 6.**
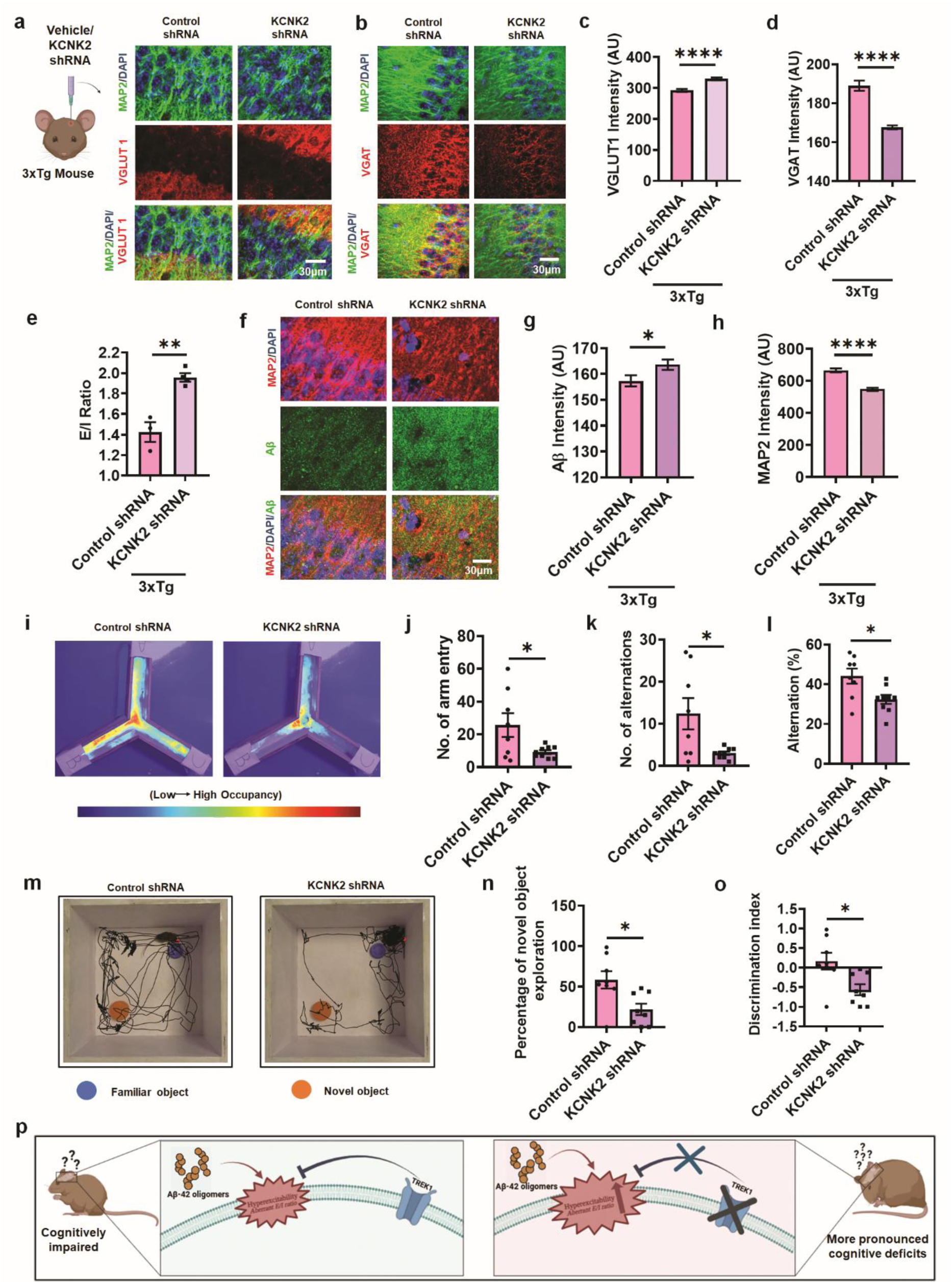
TREK1 knockdown disrupts synaptic excitatory/inhibitory balance, enhances amyloid-β deposition, and worsens dendritic integrity in 3xTg mice. a, Representative immunofluorescence images showing increased VGLUT1 intensity in the hippocampus of 3xTg mice 15 days after intrahippocampal injection with TREK1 shRNA lentivirus compared with vehicle-injected controls. b, Representative images showing decreased VGAT intensity under the same TREK1 knockdown conditions. c, Quantification of VGLUT1 fluorescence intensity in 3xTg mice after TREK1 knockdown (n = 67–76 sections; ****p < 0.0001; unpaired t-test). d, Quantification of VGAT fluorescence intensity in 3xTg mice after TREK1 knockdown (n = 60–63 sections; ****p < 0.0001; unpaired t-test). e, Quantification of Excitatory/inhibitory (E/I) ratio, calculated as VGLUT1/VGAT intensity, is markedly elevated in TREK1 knockdown mice compared with vehicle injected controls (n = 3–4; **p < 0.01; unpaired t-test). f, Representative immunofluorescence images demonstrating enhanced Aβ deposition in the hippocampus of TREK1 knockdown 3xTg mice compared with vehicle-injected controls. g, Quantification of Aβ fluorescence intensity in 3xTg mice after TREK1 knockdown (n = 50 sections; *p < 0.05; unpaired t-test). h, Quantification showing a significant decrease in MAP2 intensity in the hippocampus of 3xTg mice injected with TREK1 shRNA compared with scrambled (Sc) shRNA controls (n = 159–166 sections; ****p < 0.0001; unpaired t-test). i, Representative heat maps of spontaneous alternation behavior in the Y-maze from 3xTg mice injected with control or KCNK2 shRNA, with warmer colors indicating higher occupancy. j–l, Y-maze performance metrics showing reduced number of arm entries (j), number of alternations (k), and percent alternation (l) in TREK1 knockdown 3XTg mice relative to controls (n=8-9 mice; *p < 0.05 for all). m, Representative tracking plots from the novel object recognition (NOR) assay in 3xTg mice injected with control or KCNK2 shRNA, indicating exploration of the familiar (blue, smaller) and novel (orange, bigger) objects. n–o, NOR performance metrics in TREK1 knockdown 3xTg mice showing significantly reduced novel object exploration time (n) and discrimination index (o) compared with control shRNA treated mice (n=8 mice, *p < 0.05). p, Schematic representation illustrating the effects of TREK1 knockdown on excitatory/inhibitory balance. Data are expressed as mean ± SEM from 3–4 mice per group for IHC studies and 8-9 mice per group for cognitive assays.

### Increased TREK1 expression decreases pathological accumulation of Aβ and is essential for cognitive function in early AD

We next examined additional pathological features of AD, specifically dystrophic neurites and Aβ accumulation, in the hippocampus of 3xTg mice following TREK1 silencing. Knockdown of TREK1 resulted in a notable increase in Aβ levels, indicating enhanced amyloid accumulation (Fig. 6f, g). It is known that AD transgenic mice and AD patient hiPSC derived neurons manifest dystrophic neurites often reflected by reduced MAP2 expression^8, 45^. Accordingly, we stained for MAP2 (a marker of dendrites) and found that its expression decreased further following TREK1 knockdown in 3xTg mice suggesting exacerbation of neurite pathology (Fig. 6h and Supplementary Figure 1f, h). This enhanced TREK1 expression could be a possible survival mechanism of the neurons to limit damage at early stages of AD.

Next, we investigated how TREK1 influences cognitive performance in an AD mouse model. Consistent with established literature ^46–48^, our initial behavioral tests confirmed that 3XTg-AD mice exhibit significant cognitive impairment compared to wild-type, WT (B6) control mice. We demonstrated the difference in cognitive function between 3XTg-AD and WT (B6) mice using two standard behavioral paradigms, the Y-maze test for short-term spatial working memory ^47,48^ and the novel object recognition (NOR) test for hippocampal-dependent learning and memory, which leverages a rodent’s innate preference for novelty ^46^ (Supplementary Fig. 8 a-g). To isolate TREK1’s role, we performed hippocampal knockdown of TREK1 in 3XTg mice using shRNA, with a scrambled shRNA-injected 3XTg group as the control. Analysis of Y-maze performance revealed that TREK1 depletion significantly exacerbated cognitive deficits. Key parameters, including the total number of arm entries, the number of spontaneous alternations between arms, and the calculated percentage of alternations, were all markedly worse in the TREK1 knockdown group compared to the scrambled shRNA control (Fig. 6 i-l). A striking finding emerged from the analysis of percent alternation, a primary index of working memory. The decrease in this metric caused by TREK1 knockdown within 3XTg mice was substantial, exceeding 10%. Importantly, this deficit was quantitatively equivalent to the impairment observed between 3XTg mice and WT B6 controls (Fig. 6l, Supplementary Fig. 8d). This indicates that the removal of TREK1 from an AD background inflicts damage to short-term spatial memory that is nearly as severe as the damage caused by the AD-related genetic mutations themselves, underscoring TREK1’s critical role in preserving this cognitive domain (Fig. 6 i-l, Supplementary Fig. 8a-d).

We extended our investigation to recognition memory using the NOR test. Mirroring the Y-maze results, TREK1 knockdown in 3XTg mice led to a significant reduction in both the percentage of time spent exploring the novel object and the resulting discrimination index when compared to control 3XTg mice (Fig. 6 m-o, Supplementary Fig. 8 e-g). Again, the magnitude of the deficit was profound. The reduction in the discrimination index due to TREK1 knockdown was similar to the reduction seen when comparing untreated 3XTg mice to WT B6 mice (Fig. 6o, Supplementary Fig. 8g). This reinforces the conclusion that the absence of TREK1 in AD precipitates extreme cognitive dysfunction.

## Discussion

Our investigation reveals a critical compensatory mechanism in early AD pathogenesis, demonstrating that neurons upregulate the two-pore domain leak potassium channel TREK1 in response to Aβ-induced hyperexcitability. We observed TREK1 upregulation triggered by Aβ deposition and neuronal hyperactivity in two well-established AD mouse models - APP/PS1 and 3xTg mice. APP/PS1 mice exhibit Aβ deposition initiating in the cortex at 6 weeks postnatal, with subsequent hippocampal accumulation by 3-4 months of age^32, 49^. Electrophysiological analyses revealed concomitant development of neuronal hyperexcitability, including enhanced intrinsic excitability and synaptic reorganization leading to elevated hippocampal activity by 6 weeks^45, 50^. The 3xTg model demonstrated more aggressive pathology, showing an 8-fold elevation in cerebrocortical Aβ42 levels versus WT by 3 months^32^, and exhibited early hyperexcitability phenotypes^51, 52^, recapitulating the epileptiform activity observed in early AD patients^1, 53^. This observation in transgenic AD mice models combined with the well-documented propensity of Aβ42o to induce neuronal hyperactivity, led us to hypothesize that Aβ42o-mediated hyperexcitability affects TREK1 expression. To test the hypothesis, we first confirmed the pathology in terms of Aβ levels via IHC and higher neuronal activity via calcium imaging in hippocampal and cortical brain sections in both the transgenic mice. Notably, both models showed significant upregulation of TREK1 expression in hippocampal and cortical regions precisely coinciding with these early pathological changes (by 3 months of age; Fig. 1). Interestingly, Aβ42o treatment alone increased TREK1 levels both in rat primary neurons and in mice hippocampus suggesting that the TREK1 upregulation in AD transgenic mice was due to the accumulation of Aβ42o. Further we observed that neuronal hyperexcitability was sufficient to upregulate TREK1. On suppressing neuronal activity in 3xTg mice, TREK1 levels decreased significantly suggesting that neuronal activity was driving TREK1 expression (Fig. 2).

The hyperactivity observed in early AD pathogenesis leads to significant increase in neuronal calcium levels. We found a decrease in TREK1 levels on chelating calcium ions using BAPTA establishing the role of intracellular calcium in Aβ42o-mediated TREK1 upregulation. This increased intracellular calcium activates specific calcium-sensitive membrane-bound isoforms of ACs (AC1, AC3, and AC8)^54^. However, these isoforms exhibit markedly different calcium sensitivities and expression patterns with AC1 demonstrating the highest calcium sensitivity compared to both AC3 and AC8^55^. The expression of AC3 is mostly found in dorsal root ganglion, olfactory sensory neurons, choroid plexus cells and astrocytes^56–58^, so we did not investigate the role of AC3 in upregulation of TREK1. As the intracellular calcium levels are known to increase manifold in AD, we expected both AC1 and AC8 to get activated ^9, 37, 50^. The regional expression pattern of AC1 and AC8 in hippocampus, neocortex, and entorhinal cortex further underscores its potential importance in AD pathophysiology ^38, 59^. Genetic evidence from AC1/AC8 double knockout mice has conclusively demonstrated that these two isoforms represent the major calcium-activated ACs in neurons^40^. Interestingly, while postmortem studies of advanced AD brains have reported decreased AC1 expression^60^, the physiological role of AC1/AC8 in early AD stages remains poorly characterized where increased calcium is a hallmark of earliest detectable pathological changes in AD progression. Indeed, silencing AC1/AC8 downregulated TREK1 expression even in the presence of Aβ42o (Fig. 3).

AC1 and AC8 serve as crucial molecular bridges between neuronal activity and intracellular signalling through their generation of cAMP^54^. The cAMP-PKA pathway has been extensively characterized in multiple cellular contexts, where it serves as a fundamental signalling module for translating extracellular stimuli into transcriptional responses^61^. In our study, we found that increasing cAMP levels using either a cell-permeable cAMP analog or the activator Forskolin led to an upregulation of TREK1 expression (Fig. 3). In non-neuronal cells, cAMP elevation and PKA activation robustly increase TREK1 expression as demonstrated in astrocytes^26^ and adrenal cortical cells^39^. Crucially, we also observed that PKA inhibitors, KT5720 and H89, were able to downregulate TREK1 expression even in the presence of Aβ42o (Fig. 4).

Protein kinase A (PKA) is a well-established regulator of gene transcription, primarily through its phosphorylation and activation of the cAMP response element-binding protein (CREB)^62^. However, our bioinformatic analysis of the TREK1 (KCNK2) gene revealed that CTCF (CCCTC-binding factor) binding motifs were significantly enriched in TREK1 regulatory regions, whereas canonical CREB-binding sites were less prominent. This observation led us to investigate whether PKA regulates TREK1 expression through CTCF. Our experiments demonstrated decreased CTCF levels in response to PKA blockers in primary neurons which is in line with previous studies^43^. Further we found that on silencing CTCF in Aβ42o treated cultures and in 3xTg mice, the TREK1 expression levels decreased showing that CTCF was playing a role in its expression (Fig. 4).

Next, we observed that this increased expression of TREK1 was helping the neurons to protect themselves from the Aβ42o-related excitotoxicity. In AD models, silencing or pharmacologically inhibiting TREK1 resulted in further depolarization of neurons and a worsening of the E/I imbalance (Fig. 5 and 6). Additionally, it led to excessive spontaneous neuronal firing manifested by an increased frequency of calcium events and more frequent spontaneous action potentials (Fig. 5). TREK1 knockdown was also causing further damage to neurons reflected in greater accumulation of Aβ and increased dystrophic neurites in AD transgenic mice (Fig. 6). In AD, neurons have been shown to employ compensatory mechanisms involving proteins like transthyretin, neprilysin and clusterin to combat Aβ toxicity by inhibiting Aβ aggregation and detoxification^63–65^. Additionally, increased E-I ratio has been shown to be a compensatory mechanism in autism spectrum disorder (ASD) where similar hyperactivity is observed in several forms of ASD^66^. Starting from the accumulation of Aβ in the brain, the first sign of the disease, to the clinical diagnosis of dementia may take up to 20 years^67^. However, the early factors which contribute to the slowing of the progression of AD has always remained unclear. We find that the upregulation of TREK1 acts as a buffer to slow down cognitive decline. This is evident from the worsening performance of 3XTg mice in two standard cognitive tests, Y-maze and NOR after knocking down TREK1 in their hippocampus (Fig. 6). This further supports the idea that TREK1 upregulation is an innate mechanism to preserve cognitive function even in the face of AD pathology.

This study is the first to show that TREK1 is upregulated by a specific, activity-dependent mechanism (Aβ42-hyperactivity-AC-cAMP-PKA-CTCF) as a direct response to protect neurons from the toxic effects of Aβ in early AD (Fig. 7). We propose that the upregulation of TREK1 acts as an endogenous neuroprotective response by dampening neuronal hyperexcitability which helps in maintaining E/I ratio and may therefore slow the progression of the disease. Consequently, the study highlights the modulation of TREK1 as a homeostatic survival mechanism which can be exploited as a promising therapeutic strategy for AD.

**Fig. 7.**
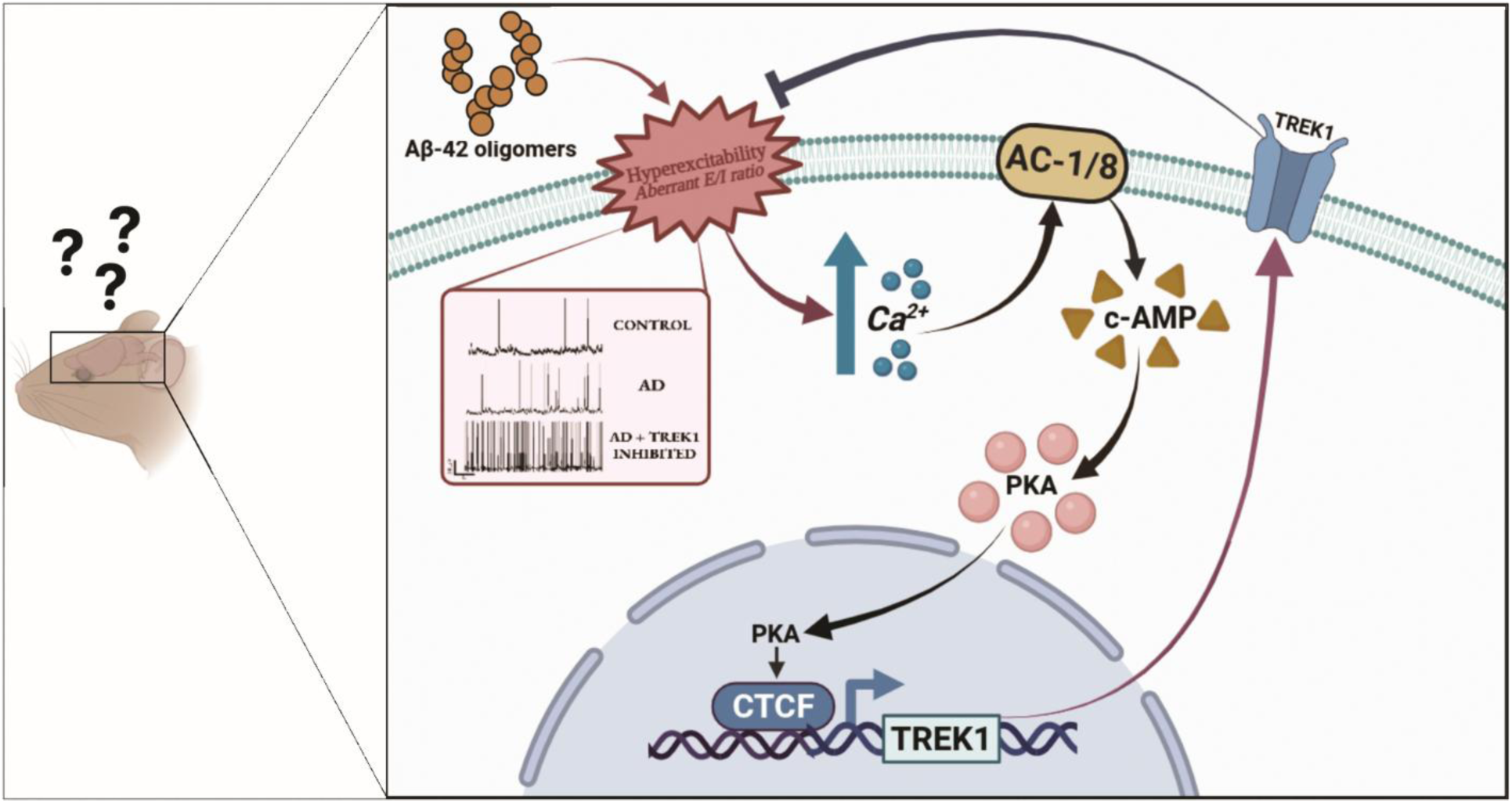
Schematic representation of the pathway underlying Aβ42 induced TREK1 upregulation and its functional consequences. The diagram summarizes the signaling cascade by which Aβ42 induced neuronal hyperexcitability drives TREK1 upregulation via calcium influx and the AC1/AC8–cAMP–PKA–CTCF axis. TREK1 upregulation decreases neuronal excitability, limits excitatory/inhibitory balance, thereby improves neuronal health in 3xTg mice.

## Methods

### Animal models

All animal experiments were conducted in accordance with CPCSEA regulations (Registration No. 1634/GO/ReRcBiBt/S/12/CPCSEA, DoR-16.05.2023) and approved by the Institutional Animal Ethics Committee (IAEC) at the National Institute of Science Education and Research (NISER), Bhubaneswar, India, and all procedures were performed following institutional and national ethical guidelines (Ethical Approval No: NISER/SBS/AH-281 and NISER/SBS/AH-280). Primary neuronal cultures were prepared from postnatal day 0-1 (P0-P1) Sprague Dawley rat pups for in vitro studies. Alzheimer’s disease mouse models included 3xTg-AD mice [B6;129-Tg(APPSwe,tauP301L)1Lfa Psen1tm1Mpm/Mmjax; MMRRC ID: 034830, JAX 004807], which develop both amyloid and tau pathology, and APP/PS1 mice [B6C3-Tg(APPswe, PSEN1dE9)85Dbo/Mmjax; MMRRC ID: 034829, JAX 004462], which exhibit early amyloid deposition, and were used to investigate in vivo disease-relevant mechanisms. C57BL/6 mice were used as WT mice for intrahippocampal injections of Aβ42o, Aβ40o and APV for deciphering the pathological mechanisms in this study.

### Primary neuron astrocyte co-culture

Cortical neurons and astrocytes were isolated from postnatal day 0–1 Sprague-Dawley rat pups. Brains were extracted and immediately transferred into ice-cold calcium- and magnesium-free Hanks’ Balanced Salt Solution (HBSS) supplemented with 10 mM glucose and HEPES (Sigma). Cortical regions were dissected and enzymatically digested using 0.25% trypsin-EDTA (Gibco) and 150 U/ml DNase I (Sigma) at 37°C for 15 minutes. Digestion was halted by adding 10% fetal bovine serum (FBS, Gibco). The tissue was then mechanically dissociated and centrifuged at 1000 rpm for 5 minutes at 4°C. The resulting cell pellet was resuspended in neurobasal-A medium (Gibco) supplemented with 10% FBS, 1% Glutamax, 1% antibiotic-antimycotic (Gibco), and 2% N2 supplement (Gibco). Cells were seeded onto coverslips pre-coated with 0.1 mg/ml Poly-D-lysine.^26^ Cultures were maintained at 37°C in a humidified incubator with 5% CO₂. The medium was replaced on day 4, and subsequently refreshed every three days. Experimental procedures were carried out on culture days 11–12.

### Preparation of Aβ oligomers

Beta-Amyloid (1-42) peptide (Anaspec) was dissolved in 1,1,1,3,3,3-hexafluoro-2-propanol (HFIP) to a final concentration of 1 mM. The solution was incubated for 2 hours at room temperature (RT). HFIP was then evaporated. The peptide film was resuspended in dimethyl sulfoxide (DMSO) to prepare a 5 mM solution, followed by bath sonication for 10 minutes. At this point, the monomeric solution can be stored at −20°C. Aβ peptides were further processed by diluting the solution in MEM to achieve a final concentration of 500 µM, followed by overnight incubation at 4°C. The prepared solution was centrifuged at 1000 rpm for 2 minutes. The supernatant containing soluble oligomers was used during treatment at a final concentration of 5 µM^68–71^.

### Transfection of primary neurons

Transfection was performed on day 9 in vitro. 500ng TREK1 expressing plasmid was used for TREK1 overexpression. Plasmid DNA and Lipofectamine 2000 (Invitrogen) were each diluted separately in Minimal Essential Medium (Opti-MEM™, Gibco) and incubated for 5 minutes at RT. The components were then combined at a 2:3 ratio (µg DNA: µL Lipofectamine) and incubated for 15 minutes to allow complex formation. Before transfection, the complete Neurobasal-A medium was removed from the cultures. The DNA–Lipofectamine complexes were added dropwise to the cells and incubated at 37°C in a 5% CO₂ incubator for 4 hours. After incubation, the transfection medium was replaced with a fresh, complete Neurobasal-A medium. Cells were maintained under standard conditions and treated 36 hours post-transfection for downstream analysis.

For siRNA transfection, Lipofectamine RNAiMAX (Invitrogen) was used along with Silencer Select siRNAs targeting ADCY1 (AC1; Assay IDs: s156880 and s156881), ADCY8 (AC8; Assay IDs: s130997 and s130998), CTCF (Assay IDs: s136223 and s136224) and KCNK2 (Assay IDs: s139431 and s 139432), as well as a Silencer Select negative control siRNA (all from Invitrogen). In each well of a 12-well plate, 20 pmol of siRNA was transfected using a 10:3 (pmol:µL) siRNA to Lipofectamine RNAiMAX ratio. siRNA and Lipofectamine RNAiMAX were diluted separately in Opti-MEM™ and incubated for 5 minutes at RT. The two components were then combined and incubated for 15 minutes to form complexes. Cells were transfected using the same procedure as plasmid transfection. After transfection, cells were maintained at 37°C in a 5% CO₂ incubator and treated 36 hours post-transfection.

### Cellular treatments

Primary neuron–astrocyte co-cultures were exposed to a series of pharmacological blockers and activators to delineate the signaling pathways mediating Aβ₄₂-induced upregulation of TREK-1 expression. Cultures were treated with Aβ₄₂ (5 μM) alone or in combination with specific inhibitors and modulators (all from Tocris Bioscience, unless otherwise indicated), including the voltage-gated sodium channel blocker tetrodotoxin (TTX, 1 μM), NMDA receptor antagonist 2-amino-5-phosphonovaleric acid (APV, 50 μM), intracellular calcium chelator BAPTA-AM (10 μM), PKA inhibitors KT5720 (5 μM) and H-89 (10 μM), adenylate cyclase inhibitor ST034307 (5 μM), TREK-1 blocker spadin (1 μM), and TREK-1 activator BL-1249 (10 μM). In parallel, to directly stimulate individual signaling nodes, cultures were treated with pathway-specific agonists in the absence of Aβ₄₂, including a glutamate:glycine mixture (100 μM glutamate from Sigma-Aldrich and 10 μM glycine; 10:1 ratio), NMDA (100 μM; Tocris), KCl (25 mM; Fisher Scientific), forskolin (5 μM; Tocris), and the cAMP analog 8-CPT-cAMP (100 μM; Santa Cruz Biotechnology). All pharmacological agents were freshly prepared in Neurobasal-A medium on the day of treatment and applied for 24 h.

### Immunocytochemistry

Following 24 hours of treatment, cells were rinsed once with 1X phosphate-buffered saline (PBS) and subsequently fixed in 4% paraformaldehyde for 30 minutes. After fixation, cells were washed three times with PBS. Blocking and permeabilization were performed using a solution of 3% bovine serum albumin (BSA) and 0.3% Triton X-100 in PBS for 45 minutes. Cells were then washed three times with PBST (PBS containing 0.1% Triton X-100). Primary antibodies- chicken MAP2 (1:1000, Invitrogen, PA1-10005), rabbit TREK1 (1:100, Alomone Labs, #APC-047), Rabbit CTCF (1:100, Invitrogen, #MA5-88115), Mouse AC1 (1:50, Santa Cruz, #SC-365350), Mouse AC8 (1:50, Santa Cruz, #SC-377442) and rabbit MAP2 (1:150, Cell Signalling Technology, #8707S) were diluted in PBST and applied overnight at 4°C. The following day, cells were washed once with PBST and twice with PBS. Secondary antibodies, Alexa Fluor 488-conjugated anti-rabbit (1:1000, Cell Signaling Technology, #4412S), Alexa Fluor 555 conjugated anti-mouse (1:1000, Cell Signalling Technology, #4409S) and Alexa Fluor 647-conjugated anti-chicken (1:1000, Invitrogen, #A21449), were diluted in PBS and incubated with the cells for 2 hours at RT in the dark. After incubation, cells were washed three times with PBS and mounted using ProLong Gold antifade reagent (Invitrogen)^8, 26^.

### Cryosectioning and immunohistochemistry

Following perfusion with 1X PBS and 4% paraformaldehyde (PFA), brains were post-fixed in 4% PFA at 4°C overnight. Tissues were then cryoprotected by sequential immersion in increasing concentrations of sucrose (15%, 20%, and 30% w/v in PBS), each for 12–24 hours at 4°C, until the brains sank. Cryoprotected brains were embedded in OCT compound (Leica) for cryosectioning. Coronal brain sections (30 μm thick) were cut using a cryostat (Leica) and mounted onto Superfrost Plus microscope slides. Sections were washed in PBS and permeabilized with 0.5% Triton X-100 in PBS for 15 minutes at RT. After blocking, sections were incubated with primary antibodies diluted in PBST for overnight at 4°C. The following primary antibodies were used: guinea pig anti-TREK1 (1:200, Alomone Labs, #APC-047-GP), rabbit anti-MAP2 (1:150, Cell Signaling Technology, #8707S), rabbit anti-CTCF (1:100, Invitrogen, #MA5-88115), rabbit anti-Aβ (1:100, Invitrogen, #44-344), guinea pig anti-Synapsin 1 (1:100, Synaptic Systems, #106 104), mouse anti-VGLUT1 (1:100, Synaptic Systems, #135 311), and mouse anti-VGAT (1:100, Synaptic Systems, #131 011). After the primary incubation, sections were washed thoroughly in PBST (3 x 5 minutes) and subsequently incubated with species-appropriate fluorophore-conjugated secondary antibodies for 2 hours at RT in the dark. The secondary antibodies used were: Alexa Fluor 488-conjugated anti-rabbit IgG (1:1000, Cell Signaling Technology, #4412S), Alexa Fluor 647-conjugated anti-guinea pig IgG (1:1000, Invitrogen, #A-21450), and Alexa Fluor 555-conjugated anti-mouse IgG (1:1000, Cell Signaling Technology, #4409S). Following secondary antibody incubation, sections were washed again in PBS and mounted using ProLong Gold antifade reagent (Invitrogen)^8, 26^.

### Western blot

Brain tissues were homogenized in RIPA lysis buffer (Sigma-Aldrich) supplemented with sodium orthovanadate (Sigma), β-glycerophosphate (MP Biomedicals), PMSF (Roche), protease inhibitor cocktail (Roche PI), and phosphatase inhibitor (Roche PhosStop) using a motorized homogenizer. The lysates were incubated on ice for 30 minutes with vortexing every 10 minutes, followed by centrifugation at 14,000 × g for 40 minutes at 4°C to collect the supernatant. Protein concentration was determined using the Bradford protein assay. Equal amounts of protein (40 µg) were mixed with 4× Laemmli buffer, boiled at 95°C for 10 minutes, and separated on 10% SDS-PAGE gels. Proteins were transferred to PVDF membranes (Millipore) using a semi-dry transfer system (Bio-Rad). Membranes were blocked with 3% BSA in TBS-T (Tris-buffered saline with 0.1% Tween-20) for 2 hours at RT, then incubated overnight at 4°C with primary antibodies: anti-rabbit TREK1 (1:300, Alomone Lab) and anti-mouse GAPDH (1:5000, Affinity), both diluted in blocking buffer. After washing, membranes were incubated with HRP-conjugated secondary antibodies (1:10,000, Invitrogen) for 2 hours at RT. Protein bands were visualized using enhanced chemiluminescence (ECL) reagents (Bio-Rad) and imaged with a chemiluminescence detection system^26^.

### Computational identification of CTCF binding sites

The TREK1 (KCNK2) gene sequence was identified, following which approximately 100 kb upstream of the transcription start site (TSS) was obtained from the UCSC Genome Browser (*hg38 assembly, GENCODE v48 annotation*)^72^. To identify potential transcription factor binding events, motif scanning was performed using MEME Suite (FIMO)^73^ with the CTCF motif from the JASPAR database (*ID: MA0139.1)*^74^. A threshold of *p <* 0.0001 was applied, and only sites located on the positive strand were used for further analysis. These predicted binding sites were then mapped back to the UCSC Genome Browser to examine their genomic context relative to TREK1 regulatory regions^72^. Promoter sequences were obtained from the Eukaryotic Promoter Database (EPDnew)^75^, enabling cross-comparison between computationally predicted CTCF motifs and experimentally known promoters. Further, we incorporated experimentally derived datasets from the ReMap atlas of ChIP-seq experiments^76^. Both the ReMap ChIP-seq track, reflecting direct experimental binding evidence, and the ReMap density track, representing aggregated signal across multiple datasets, were overlapped. This analysis revealed a CTCF binding site with higher density (ENCODE accession: EH38E1421793) located close to the TREK1 TSS. The site is supported by a ReMap density signal of ∼543 and overlaps with the predicted CTCF motif ^77^.

### Stereotaxic surgery and brain injections

Mice were anesthetized with isoflurane (5% for induction, 2.5% for maintenance). The surgical area was shaved and disinfected, and a midline scalp incision was made to expose the skull. Coordinates relative to bregma were determined using vernier calipers (posterior 1.5 mm, lateral 1.8 mm, and depth 2.5 mm)^78^, and a burr hole was drilled at the target site. A Hamilton syringe was used to deliver PBS, Aβ₄₂ (50 µM), or Aβ₄₂ (50 µM) co-administered with APV (2.5 mM) into the brain of C57BL/6 mice, and the needle was left in place for 2 min post-injection to minimize backflow. Following injection, the scalp was sutured and the animal was allowed to recover in a warmed chamber with postoperative monitoring and analgesic administration. Brains were isolated after 24 hrs of injection. For lentiviral-mediated knockdown studies, 1 × 10⁶ IFU of CTCF shRNA lentiviral particles (Santa Cruz Biotechnology, #sc-35125-V) or KCNK2 shRNA lentiviral particles (Santa Cruz Biotechnology, #sc-37181-V) or control shRNA lentiviral particles (Santa Cruz Biotechnology, #sc-108080) were stereotaxically delivered into the hippocampus of 3xTg mice in a total injection volume of 5 µL, and animals were maintained for 15 days post-injection before brain isolation.

### Calcium imaging in primary neuron-astrocyte co-culture

For the calcium assay, the loading solution was prepared by adding 10 mL of Fluo-4 Direct™ calcium assay buffer and 200 μL of 250 mM probenecid stock solution to one bottle of 2X Fluo-4 Direct™ calcium reagent (Invitrogen, #F10471). The solution was mixed 1:1 with culture media to achieve a final 1X Fluo-4 Direct™ calcium reagent concentration. After 24 hours of treatment, the co-cultures were loaded with the 1X Fluo-4 Direct™ calcium reagent. The reagent was added to the glass-bottom dish containing cells, and the cells were incubated with the solution for 30 minutes at 37°C. After incubation, the cells were incubated at RT for an additional 30 minutes. The dye was removed, and assay buffer was added during recording. Fluorescence intensity was recorded at 33 frames per second (fps) for 2 minutes^79^.

### Membrane potential imaging in primary neuron-astrocyte co-culture

Membrane potential changes were monitored using the FluoVolt Membrane Potential Kit (Invitrogen, #F10488). Primary neuron–astrocyte co-cultures were washed once with external solution before dye loading. Cells were incubated with FluoVolt dye solution (prepared by diluting the FluoVolt dye (1:1000) and PowerLoad (1:100) concentrate in the external solution) for 30 min at 37 °C in the dark. Following incubation, cells were washed gently with external solution to remove excess dye and background suppressor (1:100) was added, immediately subjected to live-cell imaging. Fluorescence recordings were performed using an epifluorescence microscope equipped with a high-speed camera (Axiocam 702 mono). Images were acquired at 33 frames/s with a 20X objective^80^.

### Whole-cell patch clamp electrophysiology

Whole-cell patch clamp recordings were performed on primary neuron–astrocyte co-cultures following 24 h of treatment with Aβ₄₂ or Aβ₄₂ in combination with spadin. Recording pipettes were fabricated from borosilicate glass with filament (Sutter Instrument) using a P-1000 puller (Sutter Instrument) to yield a resistance of 3–5 MΩ when filled with internal solution. The pipette internal solution contained (in mM): 120 K-gluconate, 5 KCl, 2 MgCl₂, 10 HEPES, 10 EGTA, and 4 Mg-ATP, adjusted to pH 7.4 with KOH and to an osmolarity of ∼290 mOsm. For external solution, Ca²⁺/Mg²⁺-free Hank’s Balanced Salt Solution (HBSS; Gibco, Gaithersburg, MD) was supplemented with 2 mM CaCl₂, 10 mM HEPES, and 20 μM glycine, adjusted to pH 7.4. All recordings were performed in the standard whole-cell configuration at room temperature using a Multiclamp 700B amplifier and a Digidata 1550B analog-to-digital converter (Molecular Devices). Data were sampled at 10 kHz, and both voltage-clamp and current-clamp protocols were applied using pCLAMP software (v11.4, Molecular Devices). Preliminary analysis and offline filtering at 500 Hz were achieved using Clampfit v11.4 (Molecular Devices). Spontaneous postsynaptic currents (sPSCs) were recorded in gap-free mode at a holding potential of −70 mV and 0 mV. Under these conditions at 25 °C, the chloride ion reversal potential was approximately −70 mV; hence, synaptic currents recorded at −70 mV represented excitatory responses. Since the cation reversal potential was approximately 0 mV, synaptic currents recorded at 0 mV were predominantly inhibitory in nature^8^. To calculate the frequency and amplitude of spontaneous synaptic events, SimplyFire software was used^81^. Spontaneous action potentials were recorded in current clamp mode. For highly depolarized neurons beyond −45mV, we injected negative current in the order of ∼20-30 pA to hyperpolarize the cells and calculate the action potential firing frequency. This was done to keep all the parameters comparable across conditions.

### Ex-vivo calcium imaging in brain slices

For live brain slice calcium imaging, high-sucrose artificial cerebrospinal fluid (ACSF) was composed of (in mM): 246 sucrose, 2 KCl, 1.25 NaH₂PO₄, 26 NaHCO₃, 10 D-glucose, 2 MgSO₄, 0.5 CaCl₂, and 1 sodium ascorbate, while maintenance ACSF contained (in mM): 125 NaCl, 3 KCl, 0.5–1 MgCl₂·6H₂O, 2 CaCl₂·2H₂O, 1.25 NaH₂PO₄, 26 NaHCO₃, and 26.5 D-glucose; all solutions were continuously bubbled with carbogen (95% O₂, 5% CO₂). Mice were anesthetized with isoflurane and decapitated, after which the brain was rapidly isolated into ice-cold oxygenated high-sucrose ACSF and horizontal slices (300 µm) were prepared on a vibratome (cutting speed: 0.08 mm/s, amplitude: 1.4 mm). Slices were transferred to oxygenated maintenance ACSF and allowed to recover at 34 °C for 15 min, followed by 15 min at RT, then incubated with Fluo-4 AM (Invitrogen, #F10471) in maintenance ACSF at 34 °C for 1 h under continuous carbogenation, and subsequently washed in fresh oxygenated maintenance ACSF for 30 min^82^. Calcium imaging was performed using an epifluorescence microscope equipped with a high-speed camera (Zeiss Axiocam 702 mono), acquiring images at 33 frames/s with a 30 ms exposure time, while slices were continuously perfused with oxygenated maintenance ACSF at RT.

### Y-maze test

Spontaneous alternation behavior was assessed in a Y-maze composed of three identical arms (30 cm length × 8 cm width × 15 cm height) arranged at 120°. Mice were habituated to the testing room for 30 min and then placed at the center of the maze and allowed to freely explore for 8 min while behavior was recorded by an overhead camera. An arm entry was considered complete when the hind paws of the mouse fully entered an arm, and the maze was cleaned with 70% ethanol between trials to eliminate olfactory cues. An alternation was defined as successive entries into three different arms in overlapping triplet sets. Spontaneous alternation (%) was calculated as the number of triads containing entries into all three arms divided by the total possible alternations, multiplied by 100. All testing was performed by experimenters blinded to genotype and treatment^83^.

### Novel Object Recognition Test

The novel object recognition task was performed in a 40 cm square open-field arena (35 cm height). Mice were first habituated to the empty arena for 8 mins for a day, followed by a familiarization session in which two identical objects were placed symmetrically and mice were allowed to explore for 8 min; exploration was defined as nose-directed investigation within ∼1.5 cm. After a 6 h retention interval, one familiar object was replaced with a novel object, and mice explored for 8 min while behavior was recorded and analyzed with automated tracking software. The discrimination index was calculated as DI=(T_novel_+T_familiar_)/(T_novel_−T_familiar_). Object identity and position were counterbalanced across animals, and all procedures were conducted under blinded conditions with arena/object cleaning (70% ethanol) between trials^84^.

### Image acquisition

High-resolution imaging of fixed samples was performed using a Leica SP8 laser-scanning confocal microscope equipped with a 63X oil-immersion objective. For each experimental condition, images were acquired using identical laser power, detector gain, and offset settings to ensure comparability of fluorescence intensity. Z-stacks were collected at 1 µm intervals, and maximum-intensity projections were generated for analysis. Images were collected by an observer blinded to experimental group.

Live-cell calcium and membrane potential imaging, as well as live brain slice calcium recordings, were conducted using a Zeiss Axio Observer epifluorescence microscope equipped with a high-speed camera (Axiocam 702 mono). Images were acquired at 33 frames s⁻¹ with a 30 ms exposure time. To minimize photobleaching and phototoxicity, excitation intensity was maintained at the lowest level sufficient for reliable signal detection.

### Data analysis

Image analysis was performed using Fiji/ImageJ (National Institutes of Health). For immunocytochemistry quantification, regions of interest (ROIs) were manually drawn identified by MAP2 staining. The mean fluorescence intensity within each ROI was measured, after subtracting background signal from adjacent cell-free areas. For immunohistochemistry, ROIs included the pyramidal layer to measure fluorescence intensity. For cortical TREK-1, Aβ, MAP2, and synaptic markers (VGLUT1, VGAT, and Synapsin 1), fluorescence was quantified across the entire field of view. For calcium and membrane potential imaging, ROIs were delineated around individual neuronal somata, and fluorescence intensity (F) was measured over time (t). Calcium transients were expressed as ΔF/F₀, where F₀ represents the mean of the ten lowest fluorescence values, and event frequency was analyzed. Electrophysiological recordings were analyzed using pCLAMP software (v11.4, Molecular Devices). Action potential frequency, excitatory postsynaptic current (EPSC) frequency and amplitude, and resting membrane potential were quantified.

### Statistical analysis

All statistical analyses were performed using GraphPad Prism 8.0.1. Data are presented as mean ± SEM from at least three independent biological replicates (n), where n represents independent cultures or animals. Comparisons between two groups were conducted using unpaired, two-tailed Student’s t-tests. For multiple-group comparisons, one-way ANOVA followed by Sidak’s test was applied as appropriate. A p-value < 0.05 was considered statistically significant. All quantitative analyses were carried out by an experimenter blinded to treatment conditions.

## Acknowledgements

SG acknowledges SERB (Grant no. SRG/2022/000117), Indian Council of Medical Research (ICMR) (Grant no. IIRP-2023-0585) and DBT (Grant no.

BT/PR48748/MED/122/329/2023) and Ben Barres Spotlight award for funding. SG also acknowledges Department of Atomic Energy (DAE) for the intra-mural financial support and for providing infrastructure facilities.TM acknowledges UGC for fellowship.

## Author contributions

Conceptualization, Investigation, Methodology: TM, RB, TC, AM, VM, HR, SG Data curation, Formal analysis, writing – original draft Preparation-TM, SG. Writing – review and editing, Funding acquisition and Supervision-SG.

## Data availability

The data supporting this study’s findings are available within the article and its Supplementary Information or from the corresponding author upon reasonable request.

## References

1. Lam AD, Deck G, Goldman A, Eskandar EN, Noebels J, Cole AJ (2017) Silent hippocampal seizures and spikes identified by foramen ovale electrodes in Alzheimer’s disease. Nat Med 23:678–680

2. Vossel KA, Beagle AJ, Rabinovici GD, et al (2013) Seizures and epileptiform activity in the early stages of Alzheimer disease. JAMA Neurol 70:1158–1166

3. Palop JJ, Mucke L (2009) Epilepsy and cognitive impairments in alzheimer disease. Arch Neurol 66:435–440

4. Palop JJ, Mucke L (2016) Network abnormalities and interneuron dysfunction in Alzheimer disease. Nat Rev Neurosci 17:777–792

5. Quiroz YT, Budson AE, Celone K, Ruiz A, Newmark R, Castrillõn G, Lopera F, Stern CE (2010) Hippocampal hyperactivation in presymptomatic familial Alzheimer’s disease. Ann Neurol 68:865–875

6. Nygaard HB, Kaufman AC, Sekine-Konno T, Huh LL, Going H, Feldman SJ, Kostylev MA, Strittmatter SM (2015) Brivaracetam, but not ethosuximide, reverses memory impairments in an Alzheimer’s disease mouse model. Alzheimers Res Ther. 10.1186/s13195-015-0110-9

7. Verret L, Mann EO, Hang GB, et al (2012) Inhibitory interneuron deficit links altered network activity and cognitive dysfunction in alzheimer model. Cell 149:708–721

8. Ghatak S, Dolatabadi N, Trudler D, Zhang X, Wu Y, Mohata M, Ambasudhan R, Talantova M, Lipton SA (2019) Mechanisms of hyperexcitability in alzheimer’s disease hiPSC-derived neurons and cerebral organoids vs. Isogenic control. Elife. 10.7554/eLife.50333

9. Busche MA, Chen X, Henning HA, Reichwald J, Staufenbiel M, Sakmann B, Konnerth A (2012) Critical role of soluble amyloid-β for early hippocampal hyperactivity in a mouse model of Alzheimer’s disease. Proc Natl Acad Sci U S A 109:8740–8745

10. Zott B, Simon MM, Hong W, Unger F, Chen-Engerer H-J, Frosch MP, Sakmann B, Walsh DM, Konnerth A A vicious cycle of b amyloid-dependent neuronal hyperactivation.

11. Hampel H, Hardy J, Blennow K, et al (2021) The Amyloid-β Pathway in Alzheimer’s Disease. Mol Psychiatry 26:5481–5503

12. Fani G, Mannini B, Vecchi G, Cascella R, Cecchi C, Dobson CM, Vendruscolo M, Chiti F (2021) Aβ Oligomers Dysregulate Calcium Homeostasis by Mechanosensitive Activation of AMPA and NMDA Receptors. ACS Chem Neurosci 12:766–781

13. Mattson MP, Cheng B, Davis D, Bryant K, Lieberburg I, Rydel RE (1992) p-Amyloid Peptides Destabilize Calcium Homeostasis and Render Human Cortical Neurons Vulnerable to Excitotoxicity.

14. Azargoonjahromi A (2024) The duality of amyloid-β: its role in normal and Alzheimer’s disease states. Mol Brain. 10.1186/s13041-024-01118-1

15. Barrow-F CJ, Yasuda A, Kenny PTM, Zagorskil MG (1992) Solution Conformations and Aggregational Properties of Synthetic Amyloid P-Peptides of Alzheimer’s Disease Analysis of Circular Dichroism Spectra.

16. Sgourakis NG, Yan Y, Mccallum S, Wang C, Garcia AE The Alzheimer’s peptides Aβ40 and 42 adopt distinct conformations in water: A combined MD / NMR study.

17. Ciccone R, Franco C, Piccialli I, Boscia F, Casamassa A, de Rosa V, Cepparulo P, Cataldi M, Annunziato L, Pannaccione A (2019) Amyloid β-Induced Upregulation of Nav1.6 Underlies Neuronal Hyperactivity in Tg2576 Alzheimer’s Disease Mouse Model. Sci Rep. 10.1038/s41598-019-50018-1

18. Jamshidi D, Watkins J, DeBoeuf K, Farley J (2023) Rapid suppression of macroscopic homomeric Kv1.2 and heteromeric Kv1.1/1.2 currents by amyloid beta (1-42) peptide: Implications for Alzheimer’s disease. Alzheimer’s & Dementia. 10.1002/alz.067946

19. Bhoi R, Mitra T, Tejaswi K, Manoj V, Ghatak S (2025) Role of Ion Channels in Alzheimer’s Disease Pathophysiology. Journal of Membrane Biology 258:187–212

20. Ulrich D (2015) Amyloid-β impairs synaptic inhibition via GABAA receptor endocytosis. Journal of Neuroscience 35:9205–9210

21. Ortiz-Sanz C, Balantzategi U, Quintela-López T, Ruiz A, Luchena C, Zuazo-Ibarra J, Capetillo-Zarate E, Matute C, Zugaza JL, Alberdi E (2022) Amyloid β / PKC-dependent alterations in NMDA receptor composition are detected in early stages of Alzheimeŕs disease. Cell Death Dis. 10.1038/s41419-022-04687-y

22. Honoré E (2007) The neuronal background K2P channels: Focus on TREK1. Nat Rev Neurosci 8:251–261

23. Ter PÉ, And E, Gá G, Czirják G, Czirják C (2010) Molecular Background of Leak K Currents: Two-Pore Domain Potassium Channels. 10.1152/physrev.00029.2009.-Two-pore

24. Ávalos Prado P, Landra-Willm A, Verkest C, Ribera A, Chassot AA, Baron A, Sandoz G (2021) TREK channel activation suppresses migraine pain phenotype. iScience. 10.1016/j.isci.2021.102961

25. Zhou M, Xu W, Liao G, Bi X, Baudry M (2009) Neuroprotection against neonatal hypoxia/ischemia-induced cerebral cell death by prevention of calpain-mediated mGluR1α truncation. Exp Neurol 218:75–82

26. Ghatak S, Banerjee A, Sikdar SK. Ischaemic concentrations of lactate increase TREK1 channel activity by interacting with a single histidine residue in the carboxy terminal domain. J Physiol. 2016 Jan 1;594(1):59–81. doi: 10.1113/JP270706. Epub 2015 Nov 17. PMID: 26445100; PMCID: PMC4704496.

27. Djillani A, Mazella J, Heurteaux C, Borsotto M (2019) Role of TREK-1 in health and disease, focus on the central nervous system. Front Pharmacol. 10.3389/fphar.2019.00379

28. Heurteaux C, Guy N, Laigle C, et al (2004) TREK-1, a K+ channel involved in neuroprotection and general anesthesia. EMBO Journal 23:2684–2695

29. Fink M, Duprat F, Lesage F, Reyes R, Romey G, Heurteaux C, Lazdunskil M (1996) Cloning, functional expression and brain localization of a novel unconventional outward rectifier K+ channel.

30. Li F, Zhou S ning, Zeng X, Li Z, Yang R, Wang X xi, Meng B, Pei W lin, Lu L (2022) Activation of the TREK-1 Potassium Channel Improved Cognitive Deficits in a Mouse Model of Alzheimer’s Disease by Modulating Glutamate Metabolism. Mol Neurobiol 59:5193–5206

31. Jankowsky JL, Slunt HH, Gonzales V, Jenkins NA, Copeland NG, Borchelt DR (2004) APP processing and amyloid deposition in mice haplo-insufficient for presenilin 1. Neurobiol Aging 25:885–892

32. Oddo S, Caccamo A, Shepherd JD, Murphy MP, Golde TE, Kayed R, Metherate R, Mattson MP, Akbari Y, Laferla FM (2003) Triple-Transgenic Model of Alzheimer’s Disease with Plaques and Tangles: Intracellular A and Synaptic Dysfunction.

33. Bobinski M, De Leon MJ, Tarnawski M, Wegiel J, Bobinski M, Reisberg B, Miller DC, Wisniewski HM (1998) Neuronal and volume loss in CA1 of the hippocampal formation uniquely predicts duration and severity of Alzheimer disease.

34. Rogers J, Morrison JH (1985) Quantitative Morphology and Regional and Laminar Distributions of Senile Plaques in Alzheimer’s Disease’.

35. Tamagnini F, Scullion S, Brown JT, Randall AD (2015) Intrinsic excitability changes induced by acute treatment of hippocampal CA1 pyramidal neurons with exogenous amyloid β peptide. Hippocampus 25:786–797

36. Ren S cheng, Chen P zhi, Jiang H hui, Mi Z, Xu F, Hu B, Zhang J, Zhu Z ru (2014) Persistent sodium currents contribute to Aβ1-42-induced hyperexcitation of hippocampal CA1 pyramidal neurons. Neurosci Lett 580:62–67

37. Ghatak S, Talantova M, Mckercher SR, Lipton SA (2021) Novel Therapeutic Approach for Excitatory/Inhibitory Imbalance in Neurodevelopmental and Neurodegenerative Diseases. Annu Rev Pharmacol Toxicol. 10.1146/annurev-pharmtox-032320

38. Masada N, Schaks S, Jackson SE, Sinz A, Cooper DMF (2012) Distinct mechanisms of calmodulin binding and regulation of adenylyl cyclases 1 and 8. Biochemistry 51:7917–7929

39. Enyeart JA, Liu H, Enyeart JJ (2010) cAMP analogs and their metabolites enhance TREK-1 mRNA and K+ current expression in adrenocortical cells. Mol Pharmacol 77:469–482

40. Wong ST, Athos J, Figueroa XA, Pineda V V, Schaefer ML, Chavkin CC, Muglia LJ, Storm DR (1999) Calcium-Stimulated Adenylyl Cyclase Activity Is Critical for Hippocampus-Dependent Long-Term Memory and Late Phase LTP.

41. Sassone-Corsi P (2012) The Cyclic AMP pathway. Cold Spring Harb Perspect Biol. 10.1101/cshperspect.a011148

42. Hirayama T, Tarusawa E, Yoshimura Y, Galjart N, Yagi T (2012) CTCF Is Required for Neural Development and Stochastic Expression of Clustered Pcdh Genes in Neurons. Cell Rep 2:345–357

43. Sen D, Maniyadath B, Chowdhury S, et al (2023) Metabolic regulation of CTCF expression and chromatin association dictates starvation response in mice and flies. iScience. 10.1016/j.isci.2023.107128

44. Palop JJ, Chin J, Roberson ED, et al (2007) Aberrant Excitatory Neuronal Activity and Compensatory Remodeling of Inhibitory Hippocampal Circuits in Mouse Models of Alzheimer’s Disease. Neuron 55:697–711

45. Šišková Z, Justus D, Kaneko H, et al (2014) Dendritic structural degeneration is functionally linked to cellular hyperexcitability in a mouse model of alzheimer’s disease. Neuron 84:1023–1033

46. Chiquita S, Ribeiro M, Castelhano J, et al (2019) A longitudinal multimodal in vivo molecular imaging study of the 3xTg-AD mouse model shows progressive early hippocampal and taurine loss. Hum Mol Genet 28:2174–2188

47. Knight EM, Martins IVA, Gümüsgöz S, Allan SM, Lawrence CB (2014) High-fat diet-induced memory impairment in triple-transgenic Alzheimer’s disease (3xTgAD) mice isindependent of changes in amyloid and tau pathology. Neurobiol Aging 35:1821–1832

48. Li JG, Chiu J, Ramanjulu M, Blass BE, Praticò D (2020) A pharmacological chaperone improves memory by reducing Aβ and tau neuropathology in a mouse model with plaques and tangles. Mol Neurodegener. 10.1186/s13024-019-0350-4

49. Garcia-Alloza M, Robbins EM, Zhang-Nunes SX, Purcell SM, Betensky RA, Raju S, Prada C, Greenberg SM, Bacskai BJ, Frosch MP (2006) Characterization of amyloid deposition in the APPswe/PS1dE9 mouse model of Alzheimer disease. Neurobiol Dis 24:516–524

50. Marc Aurel Busche et al.,Clusters of Hyperactive Neurons Near Amyloid Plaques in a Mouse Model of Alzheimer’s Disease. Science 321,1686–1689(2008). DOI:10.1126/science.1162844

51. Davis KE, Fox S, Gigg J (2014) Increased hippocampal excitability in the 3xTgAD mouse model for Alzheimer’s disease in vivo. PLoS One. 10.1371/journal.pone.0091203

52. Minkeviciene R, Rheims S, Dobszay MB, et al (2009) Amyloid β-induced neuronal hyperexcitability triggers progressive epilepsy. Journal of Neuroscience 29:3453–3462

53. Palop JJ, Chin J, Roberson ED, et al (2007) Aberrant Excitatory Neuronal Activity and Compensatory Remodeling of Inhibitory Hippocampal Circuits in Mouse Models of Alzheimer’s Disease. Neuron 55:697–711

54. Cooper DMF Molecular and cellular requirements for the regulation of adenylate cyclases by calcium.

55. Ferguson GD, Storm DR (2004) Why calcium-stimulated adenylyl cyclases? Physiology 19:271–276

56. Bishop GA, Berbari NF, Lewis J, Mykytyn K (2007) Type III adenylyl cyclase localizes to primary cilia throughout the adult mouse brain. Journal of Comparative Neurology 505:562–571

57. Devasani K, Yao Y (2022) Expression and functions of adenylyl cyclases in the CNS. Fluids Barriers CNS. 10.1186/s12987-022-00322-2

58. Hu ML, Zhang WW, Cao H, Zhang YQ (2019) Expression pattern of type 3 adenylyl cyclase in rodent dorsal root ganglion and its primary afferent terminals. Neurosci Lett 692:16–22

59. Xia Z, Choi E-J, Wang F, Blazynski C, Storm DR Type I Calmodulin-Sensitive Adenylyl Cyclase Is Neural Specific.

60. Xia Z, Refsdal CD, Merchant KM, Dorsa DM, Storm DR (1991) Distribution of mRNA for the Calmodulin-Sensitive Adenylate Cyclase in Rat Brain: Expression in Areas Assbciated with learning and Memory.

61. Kandel ER (2012) The molecular biology of memory: CAMP, PKA, CRE, CREB-1, CREB-2, and CPEB. Mol Brain. 10.1186/1756-6606-5-14

62. Lonze BE, Ginty DD (2002) Review Function and Regulation of CREB Family Transcription Factors in the Nervous System. Shaywitz and Greenberg

63. Alemi M, Gaiteiro C, Ribeiro CA, et al (2016) Transthyretin participates in beta-amyloid transport from the brain to the liver-involvement of the low-density lipoprotein receptor-related protein 1? Sci Rep. 10.1038/srep20164

64. El-Amouri SS, Zhu H, Yu J, Gage FH, Verma IM, Kindy MS, Johnson RH Neprilysin protects neurons against Aβ peptide toxicity.

65. Beeg M, Stravalaci M, Romeo M, Carrá AD, Cagnotto A, Rossi A, Diomede L, Salmona M, Gobbi XM (2016) Clusterin binds to Aβ1-42 Oligomers with high affinity and interferes with peptide aggregation by inhibiting primary and secondary nucleation. Journal of Biological Chemistry 291:6958–6966

66. Antoine MW, Langberg T, Schnepel P, Feldman DE (2019) Increased Excitation-Inhibition Ratio Stabilizes Synapse and Circuit Excitability in Four Autism Mouse Models. Neuron 101:648–661.e4

67. Jack CR, Andrews JS, Beach TG, et al (2024) Revised criteria for diagnosis and staging of Alzheimer’s disease: Alzheimer’s Association Workgroup. Alzheimer’s and Dementia 20:5143–5169

68. Chromy BA, Nowak RJ, Lambert MP, et al (2003) Self-Assembly of Aβ1-42 into Globular Neurotoxins. Biochemistry 42:12749–12760

69. Laurén J, Gimbel DA, Nygaard HB, Gilbert JW, Strittmatter SM (2009) Cellular prion protein mediates impairment of synaptic plasticity by amyloid-Β oligomers. Nature 457:1128–1132

70. Chang L, Cui W, Yang Y, et al (2015) Protection against β-amyloid-induced synaptic and memory impairments via altering β-amyloid assembly by bis(heptyl)-cognitin. Sci Rep. 10.1038/srep10256

71. Karthick C, Nithiyanandan S, Essa MM, Guillemin GJ, Jayachandran SK, Anusuyadevi M (2019) Time-dependent effect of oligomeric amyloid-β (1–42)-induced hippocampal neurodegeneration in rat model of Alzheimer’s disease. Neurol Res 41:139–150

72. James Kent W, Sugnet CW, Furey TS, Roskin KM, Pringle TH, Zahler AM, Haussler D (2002) The human genome browser at UCSC. Genome Res 12:996–1006

73. Grant CE, Bailey TL, Noble WS (2011) FIMO: Scanning for occurrences of a given motif. Bioinformatics 27:1017–1018

74. Fornes O, Castro-Mondragon JA, Khan A, et al (2020) JASPAR 2020: Update of the open-Access database of transcription factor binding profiles. Nucleic Acids Res 48:D87–D92

75. Meylan P, Dreos R, Ambrosini G, Groux R, Bucher P (2020) EPD in 2020: Enhanced data visualization and extension to ncRNA promoters. Nucleic Acids Res 48:D65–D69

76. Chèneby J, Gheorghe M, Artufel M, Mathelier A, Ballester B (2018) ReMap 2018: An updated atlas of regulatory regions from an integrative analysis of DNA-binding ChIP-seq experiments. Nucleic Acids Res 46:D267–D275

77. Abascal F, Acosta R, Addleman NJ, et al (2020) Expanded encyclopaedias of DNA elements in the human and mouse genomes. Nature 583:699–710

78. Franklin K. B. J., Paxinos G. (2008). The Mouse Brain in Stereotaxic Coordinates. New York, NY: Academic Press.

79. Ghatak S, Dolatabadi N, Gao R, et al (2021) NitroSynapsin ameliorates hypersynchronous neural network activity in Alzheimer hiPSC models. Mol Psychiatry 26:5751–5765

80. Pakhomov AG, Semenov I, Casciola M, Xiao S (2017) Neuronal excitation and permeabilization by 200-ns pulsed electric field: An optical membrane potential study with FluoVolt dye. Biochim Biophys Acta Biomembr 1859:1273–1281

81. Mori M, Rosko A, Farnsworth J, Carrasco G, Broomandkhoshbacht P, Pareja-Navarro K, Haghighi AP (2024) SimplyFire: An Open-Source, Customizable Software Application for the Analysis of Synaptic Events. eNeuro. 10.1523/ENEURO.0326-23.2023

82. Dasgupta D, Sikdar SK (2015) Calcium permeable AMPA receptor-dependent long lasting plasticity of intrinsic excitability in fast spiking interneurons of the dentate gyrus decreases inhibition in the granule cell layer. Hippocampus 25:269–285

83. Kim J, Kang H, Lee YB, Lee B, Lee D (2023) A quantitative analysis of spontaneous alternation behaviors on a Y-maze reveals adverse effects of acute social isolation on spatial working memory. Sci Rep. 10.1038/s41598-023-41996-4

84. Liu Y, Chu JMT, Yan T, Zhang Y, Chen Y, Chang RCC, Wong GTC (2020) Short-term resistance exercise inhibits neuroinflammation and attenuates neuropathological changes in 3xTg Alzheimer’s disease mice. J Neuroinflammation. 10.1186/s12974-019-1653-7

