## Supplementary Figures and Legends for "TREK1 upregulation is an endogenous mechanism delaying cognitive decline in Alzheimer’s Disease"

### Supplementary figures, figure legends, video legends

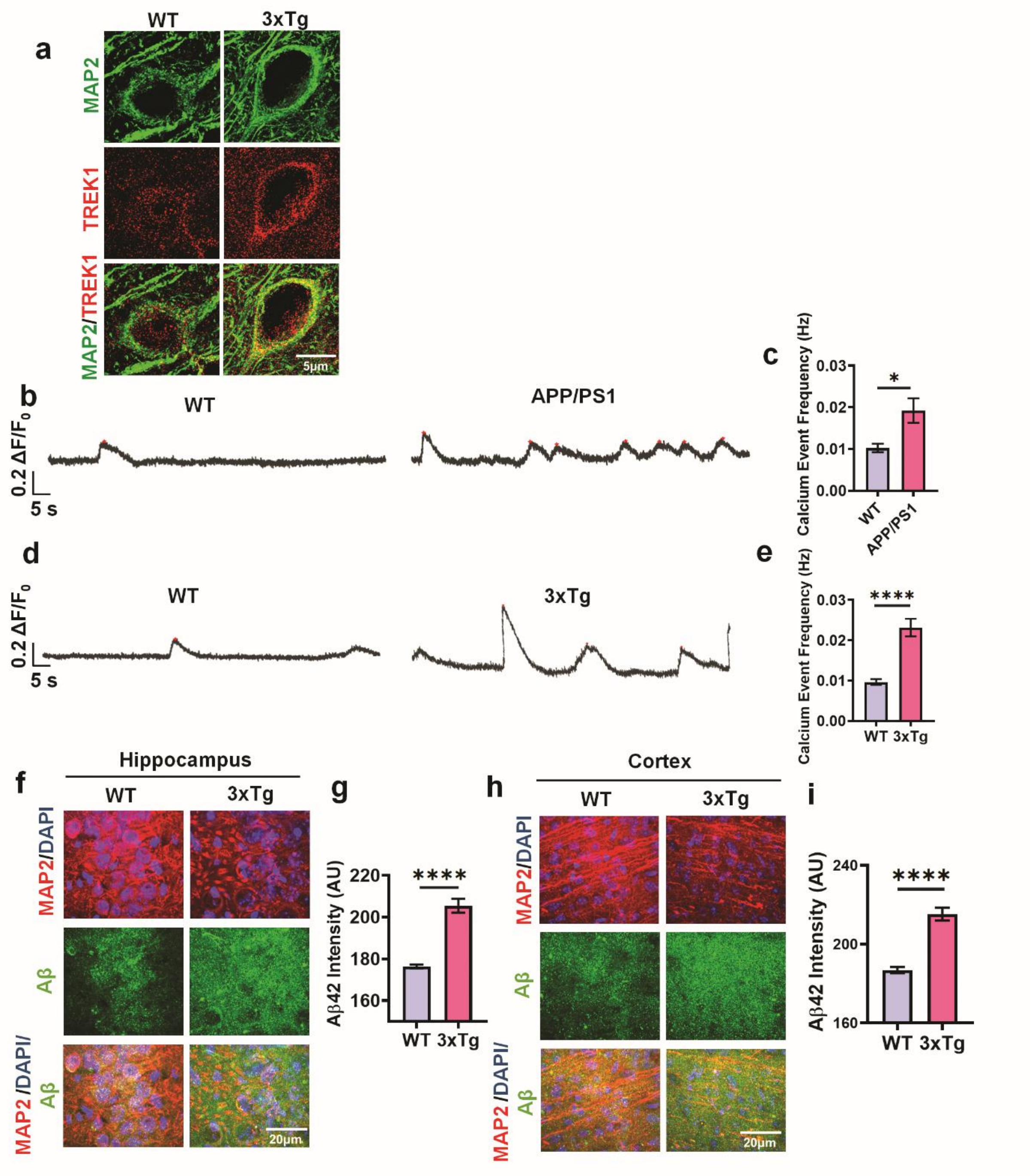


**Supplementary Fig. 1 | Increased TREK1 expression, dysregulated calcium signaling, and amyloid deposition in AD transgenic mice.**

a, Super-resolution microscopy shows elevated TREK1 expression in hippocampal neurons of 3xTg mice compared with wild-type (WT) controls.
b, Ex vivo calcium imaging of hippocampal brain sections from APP/PS1 mice reveals an increased frequency of spontaneous calcium transients relative to WT mice.
c, Quantification of calcium event frequency in hippocampal brain sections from APP/PS1 and WT mice (n=13-26 cells; $^{*}P < 0.05$; two-tailed unpaired t-test).
d, Ex vivo calcium imaging of hippocampal sections from 3xTg mice demonstrates elevated calcium transient frequency compared to WT mice.
e, Quantification of calcium event frequency in hippocampal brain sections from 3xTg and WT mice.(n=19-27 cells; $^{****}P < 0.0001$; two-tailed unpaired t-test).
f, Representative hippocampal images show markedly increased Aβ deposition in 3xTg mice compared with WT controls.
g, Quantification of Aβ fluorescence intensity in the hippocampus of 3xTg and WT mice (n=39-43 sections; $^{****}P < 0.0001$; two-tailed unpaired t-test).
h, Representative cortical images show enhanced Aβ42o deposition in 3xTg mice compared with WT controls.
i, , Quantification of Aβ fluorescence intensity in the cortex of 3xTg and WT mice (n=36-37 sections; $^{****}P < 0.0001$; two-tailed unpaired t-test).

Data are presented as mean ± SEM. n = 3-5 independent cultures or animals per group.

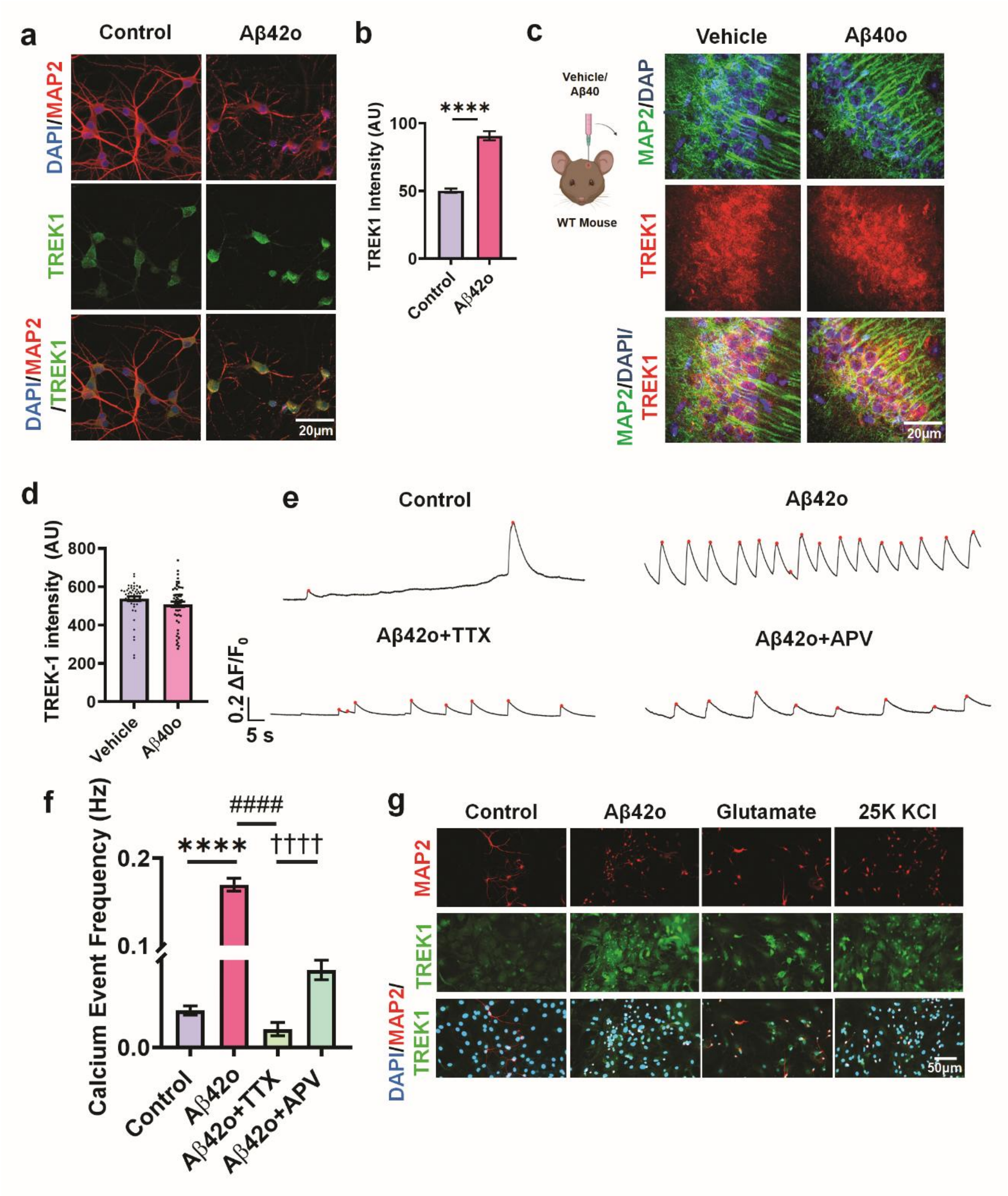


**Supplementary Fig. 2 | TREK1 upregulation under Aβ42-induced hyperexcitability.**

a, TREK1 expression is increased in hippocampal primary neuron–astrocyte cocultures following treatment with 5 μM Aβ42o.
b, Quantification of TREK1 fluorescence intensity upon treatment with Aβ42o ($n = 32$–$60$ cells; ****$P < 0.0001$; two-tailed unpaired t-test).
c, Stereotaxic intrahippocampal injection of $A\beta_{40}$o does not alter TREK1 expression.
d, Quantification of TREK1 fluorescence intensity upon $A\beta_{40}$o/vehicle injection ($n = 54$–$58$ sections).

e, Primary neuronal cultures treated with Aβ42o exhibit enhanced hyperexcitability, which is attenuated upon co-treatment with the voltage-gated sodium channel blocker TTX or the NMDA receptor antagonist APV.
f, Quantification of calcium event frequency upon treatment with Aβ42o/ Aβ42o + APV/ Aβ42o + TTX (n = 93–161 cells; ****$P < 0.0001$, ####$P < 0.0001$, ††††$P < 0.0001$; one-way ANOVA with Šidák's test).
g, Representative field images showing elevated TREK1 expression in response to hyperexcitability induced by glutamate or KCl stimulation.

Data are presented as mean ± SEM. n = 3-5 independent cultures or animals per group.

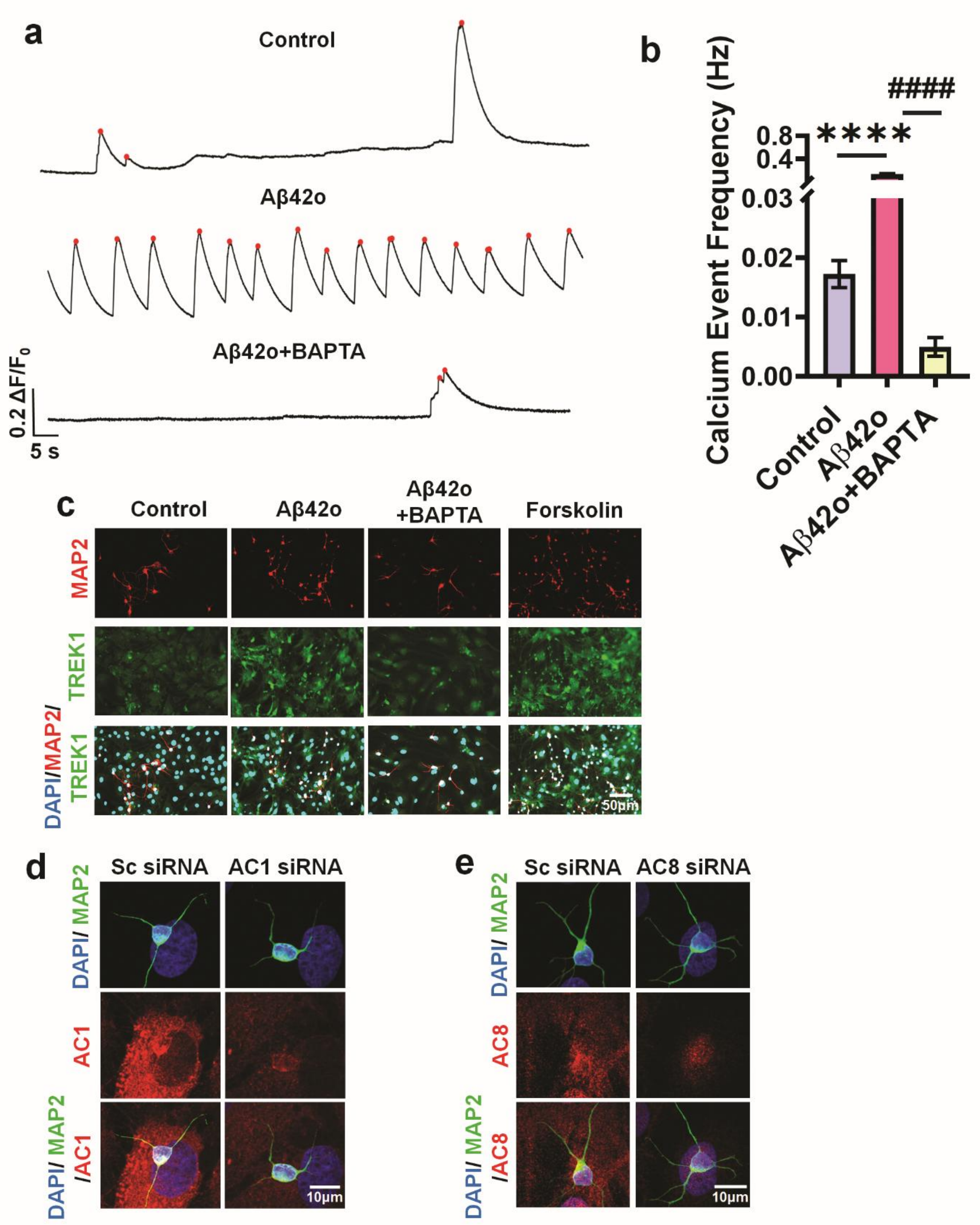


**Supplementary Fig. 3 | Calcium-dependent regulation of TREK1 expression via the cAMP pathway under hyperexcitable conditions.**

a, Representative calcium traces show that BAPTA-AM, a membrane-permeable calcium chelator, effectively reduces Aβ42o-induced neuronal hyperexcitability.
b, Quantification of calcium event following Aβ42o and/or BAPTA-AM treatment (n = 135-215 cells; ****P < 0.0001, ####P < 0.0001, ††††P < 0.0001; one-way ANOVA with Šidák's test).
c, Representative field images show that TREK1 expression fails to increase in

Aβ42o treated neurons when intracellular calcium is buffered by BAPTA-AM. In contrast, forskolin, a cAMP pathway activator, enhances TREK1 expression even in the absence of Aβ42o.
d, Primary neuron astrocyte co-cultures treated with AC1 siRNA show reduced AC1 expression, confirmed by AC1 immunostaining.
e, Primary neuron astrocyte co-cultures treated with AC8 siRNA show decreased AC8 expression, confirmed by AC8 immunostaining.

Data are presented as mean ± SEM. n = 3-5 independent cultures per group.

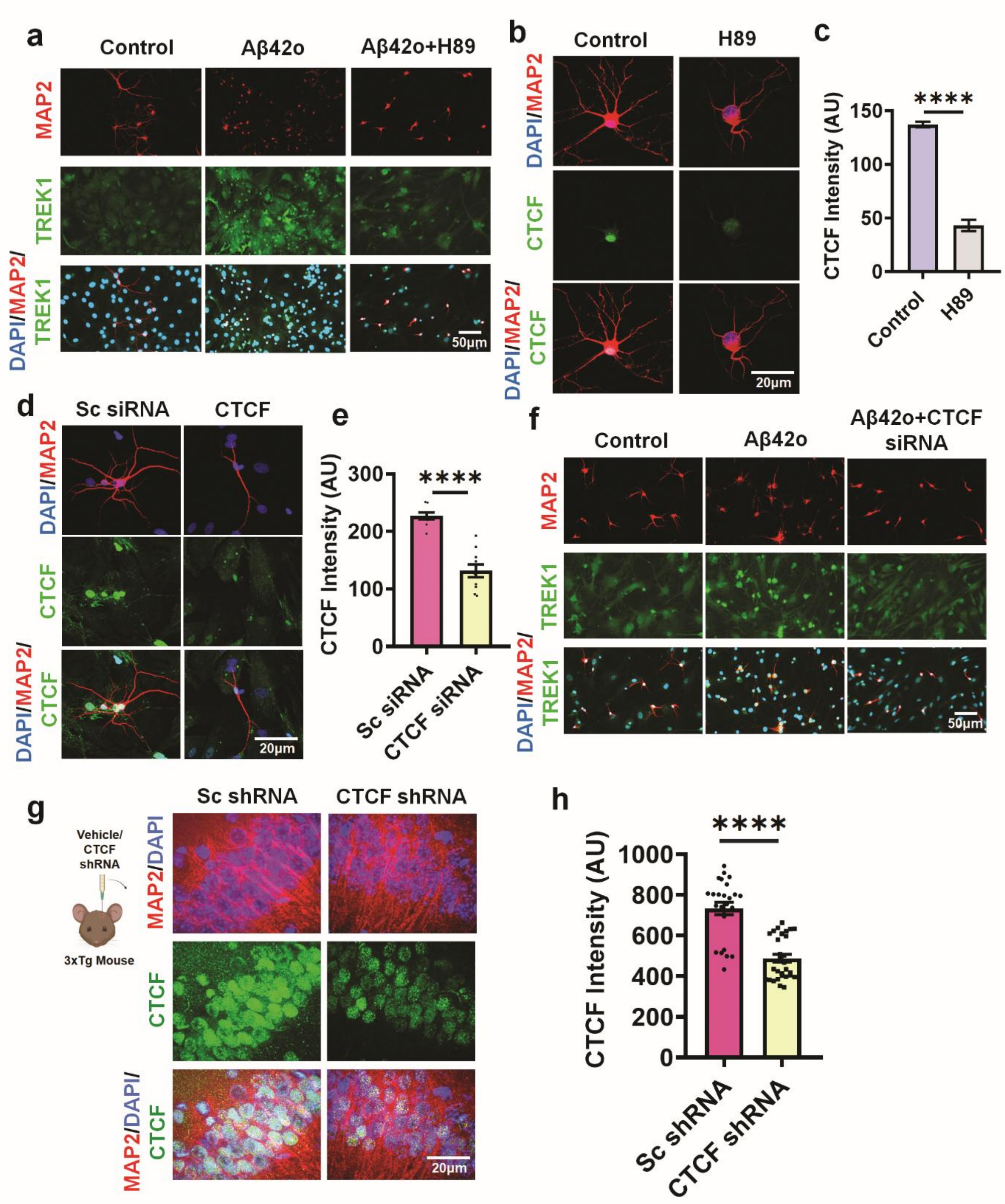


**Supplementary Fig. 4 | Regulation of TREK1 expression via the PKA–CTCF signaling pathway.**

a, Representative field images of primary neuron astrocyte co-cultures treated with Aβ42o in the presence of the PKA inhibitor H89 show reduced TREK1 expression compared with Aβ42o treatment alone.
b, PKA inhibition downregulates CTCF expression; treatment with H89 alone decreases CTCF levels.

c, Quantification of CTCF fluorescence intensity upon H89 treatment (n = 59-109 cells; ****P < 0.0001; two-tailed unpaired t-test).
d, Primary neuron-astrocyte co-cultures treated with CTCF siRNA show decreased CTCF expression, validated by CTCF immunostaining.
e, Quantification of CTCF fluorescence intensity following CTCF siRNA treatment (n = 9-10 cells; ****P < 0.0001; two-tailed unpaired t-test).
f, Field images of TREK1 immunostaining in CTCF siRNA treated cultures exposed to Aβ42o demonstrate reduced TREK1 levels compared with Aβ42o treatment alone.
g, Stereotaxic injection of CTCF shRNA lentiviral particles into the hippocampus of 3xTg mice reduces CTCF expression as compared with Sc shRNA injected mice.
h, Quantification of hippocampal CTCF fluorescence intensity in 3xTg mice after CTCF knockdown (n = 24-31 sections; ****P < 0.0001; two-tailed unpaired t-test).

Data are presented as mean ± SEM. n = 3-5 independent cultures or animals per group.

a

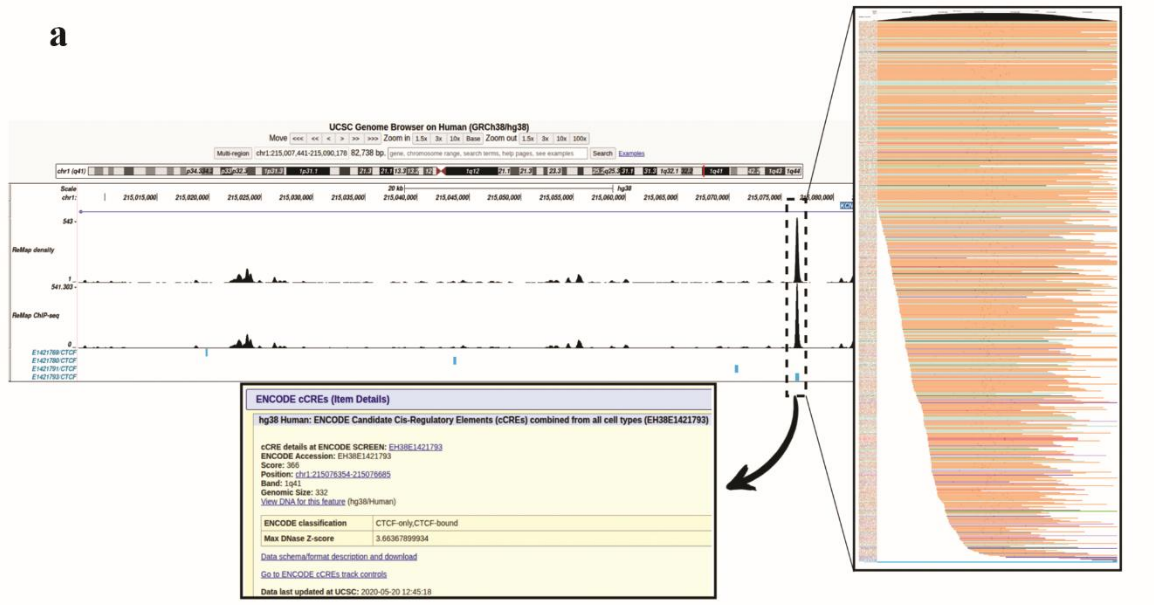


**Supplementary Fig. 5 | CTCF occupancy at the KCNK2 (TREK1) locus.**

a, UCSC Genome Browser (hg38) snapshot of the upstream KCNK2 locus (chr1:215,007,441–215,090,178) displaying ReMap density and ChIP-seq tracks. Four ENCODE-annotated CTCF sites (E1421769, E1421780, E1421791, and E1421793) are indicated, with E1421793 showing the strongest enrichment signal near the TREK1 TSS. ENCODE metadata (accession EH38E1421793; score = 366; genomic size = 332 bp; classification = CTCF-only/CTCF-bound) provide further validation. The right panel shows aggregated ChIP-seq signal across multiple cell types, confirming robust and reproducible CTCF occupancy at this site.

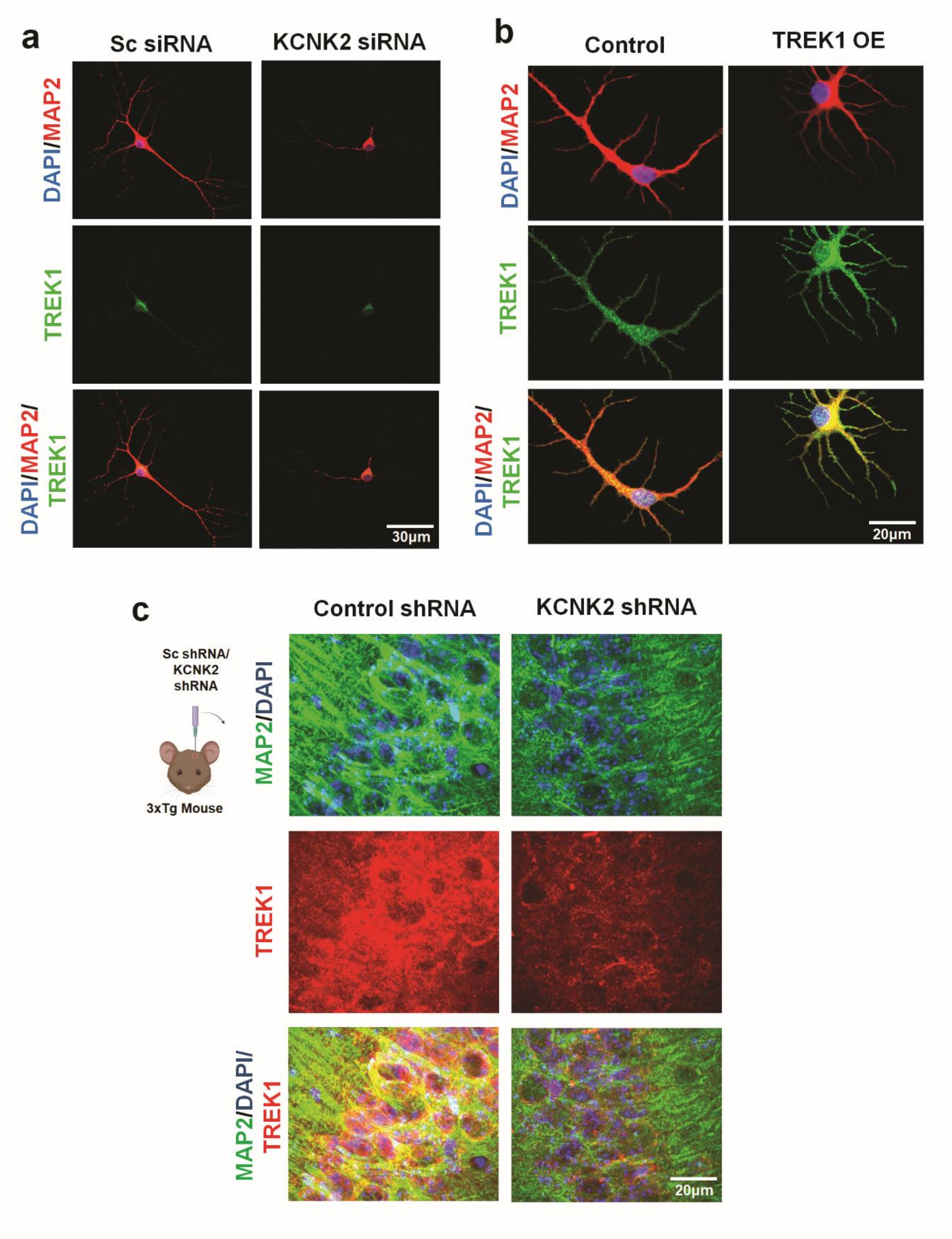


**Supplementary Fig. 6 | Modulation of TREK1 expression in vitro and in vivo.**

a, TREK1 siRNA effectively reduces TREK1 expression in primary neuronal cultures. Representative immunofluorescence images show decreased TREK1 fluorescence intensity in siRNA-treated neurons compared with control.
b, Overexpression (OE) of TREK1 in primary neurons increases TREK1 levels,

confirming successful upregulation.
c, Stereotaxic injection of TREK1 shRNA-expressing lentiviral particles into the hippocampus of mice reduces TREK1 expression in vivo. Representative hippocampal sections show diminished TREK1 immunoreactivity compared with vehicle-injected animals.

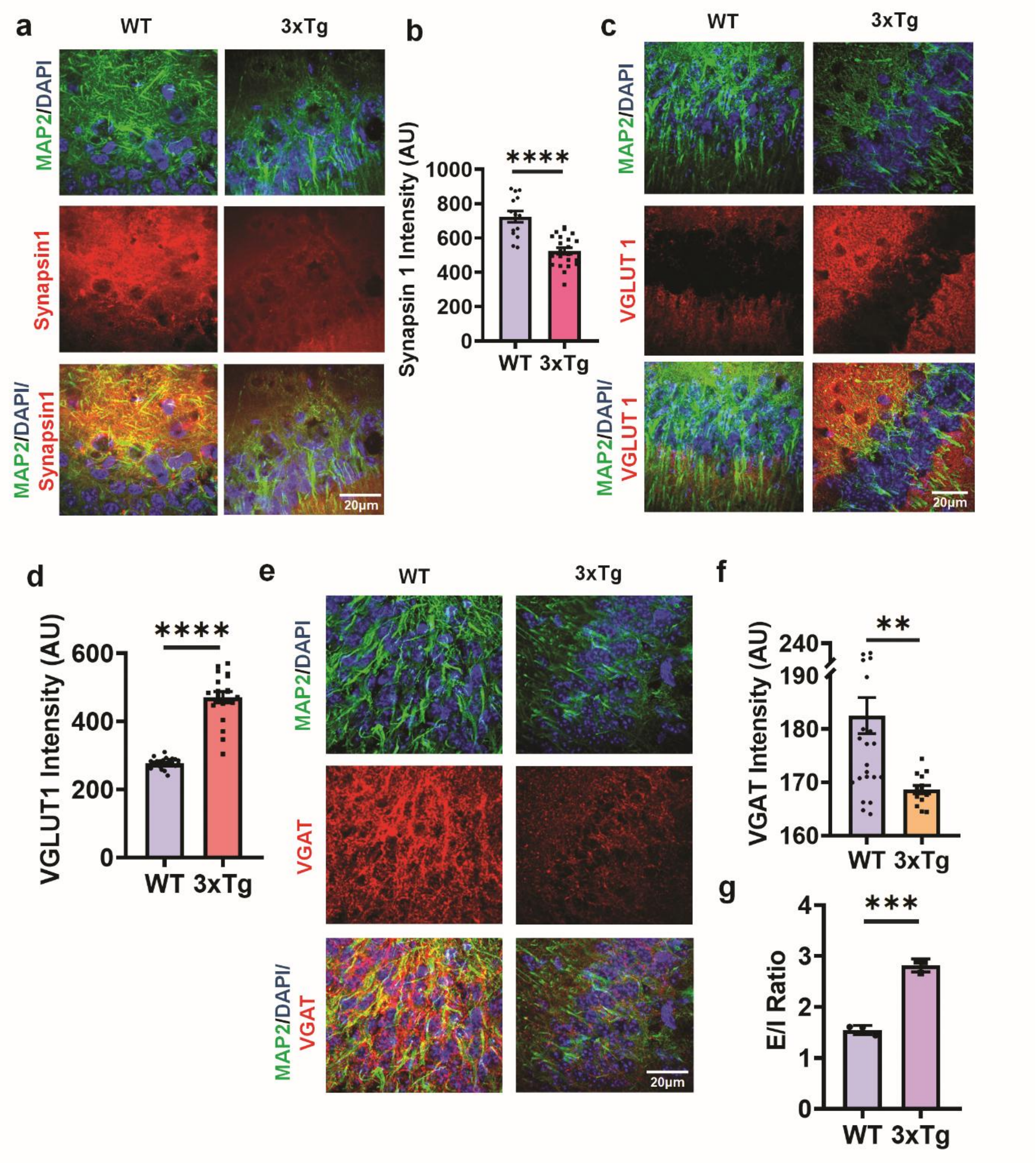


**Supplementary Fig. 7 | Synaptic alterations and increased excitatory/inhibitory (E/I) ratio in 3xTg mice.**

a, Synapsin-1, a marker of total synapses, is markedly decreased in 3xTg mice as compared to WT mice, indicating synaptic loss.

b, Quantification of Synapsin-1 fluorescence intensity in 3xTg mice compared with wild-type controls (n = 14-22 sections; ****$P < 0.0001$; two-tailed unpaired t-test).
c, VGLUT1, a presynaptic marker of excitatory glutamatergic synapses, is significantly increased in 3xTg mice compared with WT, suggesting enhanced excitatory input.
d, Quantification of VGLUT1 fluorescence intensity in 3xTg mice compared with WT controls (n = 19-26 sections; ****$P < 0.0001$; two-tailed unpaired t-test).
e, VGAT, a presynaptic marker of inhibitory GABAergic synapses, is reduced in 3xTg mice, indicating impaired inhibitory transmission.
f, Quantification of VGAT fluorescence intensity in 3xTg mice compared with WT controls (n = 14-24 sections; ****$P < 0.0001$; two-tailed unpaired t-test).
g, The resulting E/I ratio is significantly elevated in 3xTg mice, reflecting a shift toward excitatory dominance and potential hippocampal hyperexcitability (n = 3; ***$P < 0.001$; two-tailed unpaired t-test).

Data are presented as mean ± SEM. n = 3-4 animals per group.

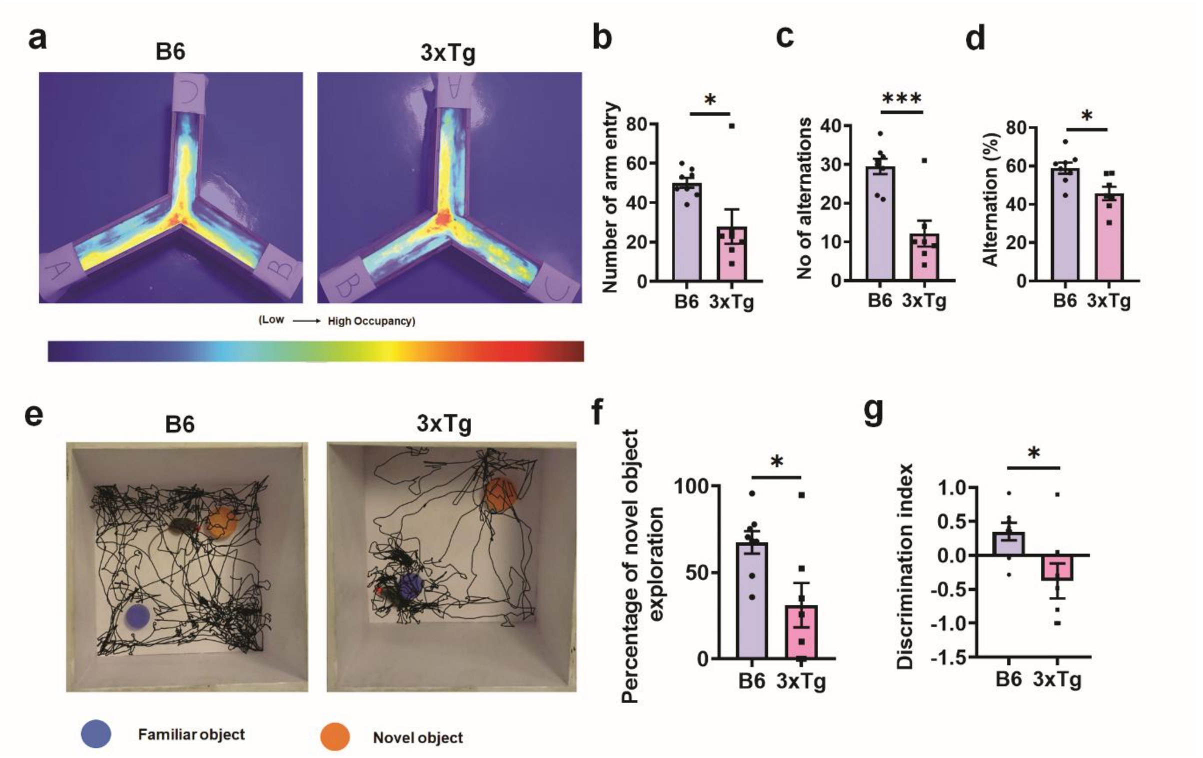


**Supplementary Fig. 8 | Impaired spatial working memory and reduced novel object recognition performance in 3xTg mice as compared with WT controls.**

**a,** Representative heatmaps of Y-maze exploration showing spatial occupancy during the 8-min session. Warmer colors indicate higher dwell time within each arm (A, B, C).

**b,** Quantification of the number of arm entries in the Y-maze, showing overall locomotor activity across groups (n = 7–8 animals per group; *P < 0.05; two-tailed unpaired t-test).

**c,** Quantification of the number of spontaneous alternations in the 3xTg mice compared with controls (n = 7–8 animals per group; ***P < 0.001; two-tailed unpaired t-test).

**d,** Quantification of the percentage of spontaneous alternations, a measure of spatial working memory, which is reduced in the 3xTg mice compared with controls (n = 7–8 animals per group; *P < 0.05; two-tailed unpaired t-test).

**e,** Representative tracking traces from the novel object recognition (NOR) test showing exploration of the familiar object (blue) and the novel object (orange) during the 8-min test session.

**f,** Quantification of the percentage of novel object exploration in the 3xTg mice compared with controls (n = 7–8 animals per group; *P < 0.05; two-tailed unpaired t-test).

**g,** Quantification of the discrimination index in the 3xTg mice compared with controls (n = 7–8 animals per group; *P < 0.05; two-tailed unpaired t-test).

Data are presented as mean ± SEM.

**Supplementary Video 1.** Representative calcium imaging showing calcium transients in control neurons exhibiting regular calcium activity. Time stamps indicate seconds.

**Supplementary Video 2.** Representative calcium imaging showing calcium transients in Aβ42o-treated neurons displaying increased calcium event frequency and hyperexcitability. Time stamps indicate seconds.

**Supplementary Video 3.** Representative calcium imaging showing calcium transients in neurons co-treated with Aβ42o and Spadin, exhibiting increased calcium event frequency compared with Aβ42o alone. Time stamps indicate seconds.

**Supplementary Video 4.** Representative calcium imaging showing calcium transients in neurons co-treated with Aβ42o and BL-1249, exhibiting decreased calcium event frequency compared with Aβ42o alone. Time stamps indicate seconds.

**Supplementary Video 5.** Representative calcium imaging showing calcium transients in neurons transfected with Sc siRNA, exhibiting regular calcium activity. Time stamps indicate seconds.

**Supplementary Video 6.** Representative calcium imaging showing calcium transients in neurons transfected with Sc siRNA and treated with Aβ42o, displaying increased calcium transients and hyperexcitability. Time stamps indicate seconds.

**Supplementary Video 7.** Representative calcium imaging showing calcium transients in neurons transfected with KCNK2 siRNA and treated with Aβ42o, exhibiting increased calcium event frequency compared with Aβ42o alone. Time stamps indicate seconds.

**Supplementary Video 8.** Representative calcium imaging showing calcium transients in neurons overexpressing KCNK2 and treated with Aβ42o, exhibiting reduced calcium event frequency compared with Aβ42o alone. Time stamps indicate seconds.

**Supplementary Video 9.** Representative ex vivo calcium imaging showing calcium transients in hippocampal sections from WT mice exhibiting regular calcium activity. Time stamps indicate seconds.

**Supplementary Video 10.** Representative ex vivo calcium imaging showing calcium transients in hippocampal sections from 3xTg mice displaying increased calcium event frequency and hyperexcitability. Time stamps indicate seconds.

**Supplementary Video 11.** Representative ex vivo calcium imaging showing calcium transients in hippocampal sections from 3xTg mice injected with Sc shRNA. Time stamps indicate seconds.

**Supplementary Video 12.** Representative ex vivo calcium imaging showing increased calcium transients in hippocampal sections from 3xTg mice injected with KCNK2 shRNA. Time stamps indicate seconds.
